# Spinal cord perfusion impairments in the M83 mouse model of Parkinson’s disease

**DOI:** 10.1101/2024.04.27.591432

**Authors:** Benjamin F. Combes, Sandeep Kumar Kalva, Pierre-Louis Benveniste, Agathe Tournant, Man Hoi Law, Joshua Newton, Maik Krüger, Rebecca Z. Weber, Inês Dias, Daniela Noain, Xose Luis Dean-Ben, Uwe Konietzko, Christian R. Baumann, Per-Göran Gillberg, Christoph Hock, Roger M. Nitsch, Julien Cohen-Adad, Daniel Razansky, Ruiqing Ni

## Abstract

Metabolism and bioenergetics in the central nervous system play important roles in the pathophysiology of Parkinson’s disease (PD). Here, we employed a multimodal imaging approach to assess oxygenation changes in the spinal cord of a transgenic M83 murine model of PD in comparison to non-transgenic littermates at 9-12 months-of-age. A lower oxygen saturation (SO_2_)^SVOT^ was detected *in vivo* with spiral volumetric optoacoustic tomography (SVOT) in the spinal cord of M83 mice compared to non-transgenic littermate mice. *Ex-vivo* high-field T1-weighted magnetic resonance imaging (MRI) and immunostaining for alpha-synuclein (phospho-S129) and vascular organisation (CD31 and GLUT1) were used to investigate the nature of the abnormalities detected via *in vivo* imaging. *Ex-vivo* analysis showed that the vascular network in the spinal cord was not impaired in the spinal cord of M83 mice. *Ex-vivo* MRI assisted with deep learning-based automatic segmentation showed no volumetric atrophy in the spinal cord of M83 mice compared to non-transgenic littermates, whereas nuclear alpha-synuclein phosphorylated at Ser129 site could be linked to early pathology and metabolic dysfunction. The proposed and validated non-invasive high-resolution imaging tool to study oxygen saturation in the spinal cord of PD mice holds promise for assessing early changes preceding motor deficits in PD mice.

## 1. Introduction

Parkinson’s disease (PD) is the second most common neurodegenerative disease affecting 1 to 3% of the elderly population (≥60 years old), with an increased prevalence over the past generation(2018). PD is defined by the appearance of clinical features, including bradykinesia, rigidity, and resting tremor (Poewe et al. 2017). The proposed new biological definition (Simuni et al. 2024) and classification of PD (Höglinger et al. 2024) suggest disease staging based on the presence of abundant intracellular aggregates termed Lewy bodies, which are mainly composed of insoluble alpha-synuclein (α-syn) fibrils that hypothesized to trigger selective and progressive neuronal death, regardless of any clinical manifestations. α-Syn undergoes posttranslational modifications, particularly phosphorylation at the Ser129 site(Anderson et al. 2006; Fujiwara et al. 2002), inducing endogenous α-syn to form these amyloid fibrils and spread within the central nervous system in a prion-like manner (Goedert et al. 2010; Hijaz and Volpicelli-Daley 2020). According to the Braak staging of the spreading of α-syn pathology, the spinal cord is affected early in the disease process, at stage 2, even prior to the nigrostriatal pathway (stage 3) and limbic system (stage 4). This is also associated with functional connectivity changes (Braak and Del Tredici 2017; Braak et al. 2003; Landelle et al. 2023), which may contribute to clinical non-motor and motor symptoms commonly developing in PD patients, including pain, constipation, and poor balance, etc (Del Tredici and Braak 2012). Recent study in animal models of PD showed that spinal cord stimulation could restores locomotion (Fuentes et al. 2009), indicating the its importance in disease progression and intervention.

Although the disease mechanisms underlying PD are still not fully understood and disease-modifying therapies still represent an unmet clinical need, growing evidence shows a tight relation between reductions in oxygen supply and α-syn pathology, PD development, and neurodegeneration (Guo et al. 2022; Lestón Pinilla et al. 2021). Indeed, the high energetic demand of neurons renders them particularly vulnerable to hypoxia. Oxygen intake and utilization disorders, such as cerebral hypoperfusion (Melzer et al. 2011) and hypoxia in the brain, have been implicated in the pathogenesis of PD (Guo et al. 2022; Pang et al. 2019). Risk genes for PD, such as leucine-rich-repeat kinase 2, ATP13A2, PTEN-induced kinase 1 and Parkin (Blauwendraat et al. 2020), are often associated with mitochondrial dysfunction and cellular respiration. In addition, environmental risk factors associated with PD, including air pollution, pesticide and heavy metal exposure, are directly linked to oxygen uptake and utilization disorders by modulating ventilation, competing for haemoglobin binding, or affecting the mitochondrial electron transport chain (Murata et al. 2022) (Hatcher et al. 2008) (Burtscher et al. 2024).

Phospho-α-syn (p-α-syn) at Ser129 site (pS129-α-syn) exists at a low level in the healthy brain and plays a physiological role at synapses (Parra-Rivas et al. 2023; Ramalingam et al. 2023). Nevertheless, Ser129 site is implicated in all pathologic α-syn aggregates, making this post-translational modification an established pathological marker of α-synucleinopathy (Anderson et al. 2006; Fujiwara et al. 2002). Recent *in vitro* studies have shown that hypoxic stress induced abnormal accumulation of pS129-α-syn, α-syn oligomer and degeneration of dopaminergic neurons (Guo et al. 2023; Li et al. 2022). Transient focal ischemia has also been shown to upregulate pS129-α-syn in a stroke mouse model (Kim et al. 2016). In humans, studies have shown increased plasma levels of total α-syn and pS129-α-syn in patients suffering from chronic intermittent hypoxia or obstructive sleep apnea (Sun et al. 2019), which are predisposing factors for PD development. Further exploration of hypoxia signalling mechanisms in α-syn pathology and neurodegeneration will facilitate a better understanding of PD pathogenesis and allow for the exploration of new therapeutic strategies.

The current study aims to develop a tool to measure *in vivo* changes in oxygenation in the spinal cord of a mouse model of α-synucleinopathy using spiral volumetric optoacoustic tomography (SVOT) (Kalva et al. 2023a, b; Ron et al. 2019b) and to assess potential structural volumetric reduction by using *ex vivo* high-field magnetic resonance imaging (MRI). A number of imaging modalities, such as perfusion and functional MRI (Laakso et al. 2021; Matsubayashi et al. 2018; Meyer et al. 2021), two-photon phosphorescence lifetime microscopy (Esipova et al. 2019; Wu et al. 2022), positron emission tomography (Mondal et al. 2021), contrast enhanced (Harmon et al. 2024) and ultrasound localization microscopy (Claron et al. 2021), have previously been employed for imaging the functional alterations in the spinal cord of rodent models. However, SVOT provides the unique combination of rapid whole-body imaging capacity (Kalva et al. 2022), high spatio-temporal resolution, and rich spectroscopic optical contrast, enabling the direct oxygenation measurements that are unachievable with other modalities (Ron et al. 2019a). We used the M83 PD model overexpressing the mutated A53T α-syn form at the early stage of pathology (Giasson et al. 2002) and assessed spinal oxygenation by mapping oxygenated hemoglobin (HbO), deoxygenated hemoglobin (HbR), and oxygen saturation (sO_2_) *in vivo* with SVOT. This was followed by *ex vivo* MRI and deep learning-based grey and white matter quantification, as well as immunofluorescence staining for validation *ex vivo* (Fig. 1).

**Figure 1.**
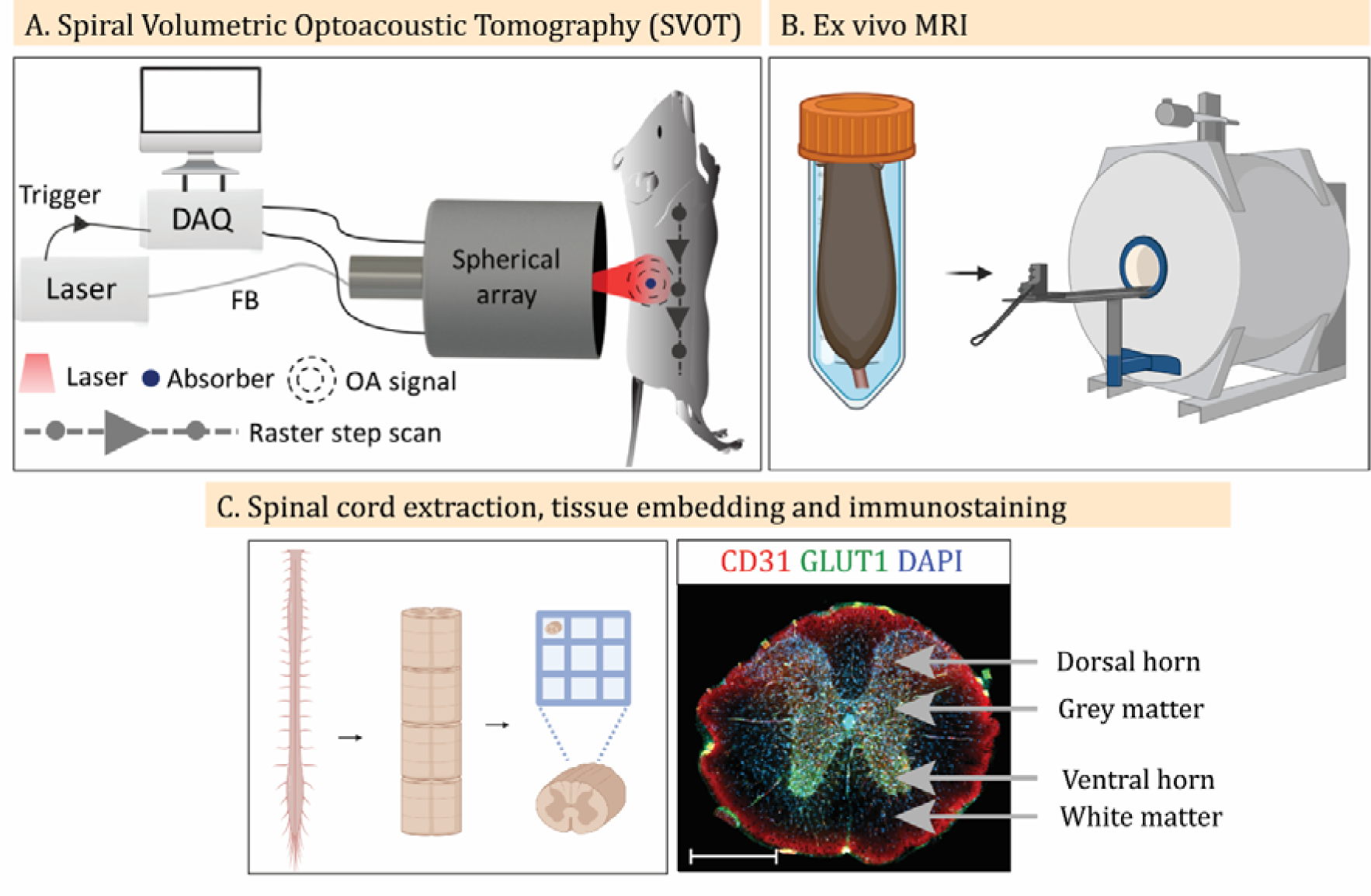
Workflow and experimental setup of the study. A) Schematic of the SVOT system for head-to-tail volumetric imaging of mice. FB: fiber bundle, DAQ: data acquisition system, OA: optoacoustic. B) *Ex vivo* MRI of the mouse spinal cord. C) Schematic of spinal cord extraction, sectioning, embedding in SpineRacks (Fiederling et al. 2021) and immunostaining. Scale bar = 50 µm. (Illustration created on Biorender.com).

## 2. Results

### 2.1 Reduction in oxygenation saturation in the spinal cord of M83 mice

We first assessed in vivo whether there was alternation in the oxygenation saturation levels in the spinal cord of asymptomatic M83 mice and NTL mice at 9-12 months of age (Fig. 1A). SVOT imaging was performed using the spherical array transducer together with the fiber bundle on the back of the mice from neck-to-tail in a step-and-go scanning manner. At each position of the spherical array, multispectral optoacoustic data was spectrally unmixed to differentiate HbO (Fig. 2A) and HbR (Fig. 2B) content along the spinal cord (Kalva et al. 2023b), followed by calculation of the SVOT-derived oxygen saturation (sO_2_ ^SVOT^) (Fig. 2C). We observed that sO_2_ ^SVOT^ levels were lower in M83 mice than in NTL mice. To visualize the differences in the sO_2_^SVOT^ across the spinal cord segments, *ex vivo* structural T1-weighted (T1w) scans were acquired to use landmarks for segmentation (Fig. 1B and Fig. 3A). sO_2_ ^SVOT^ sagittal maximum intensity projections were segmented into thoracic, lumbar, and sacral regions of interest (Fig. 3B and 3C), and the mean sO_2_^SVOT^ was computed (Fig. 3D). The cervical part of the spinal cord could not be assessed as too little optoacoustic signal was measured. This may be due to the brown adipose tissue covering this region that is highly absorbing, preventing the light from reaching the spinal cord underneath. Compared to age-matched NTL mice, M83 mice exhibited significant reductions in the sO_2_^SVOT^ in the thoracic (by 15%, p=0.0013), lumbar (by 13%, p=0.0015), sacral regions (by 16%, p=0.0005) as well as total spinal cord (by 14%, p=0.0014) (Fig. 3D). This suggested a reduction in sO_2_ level in the spinal cord of M83 mice preceding motor symptoms.

**Figure 2.**
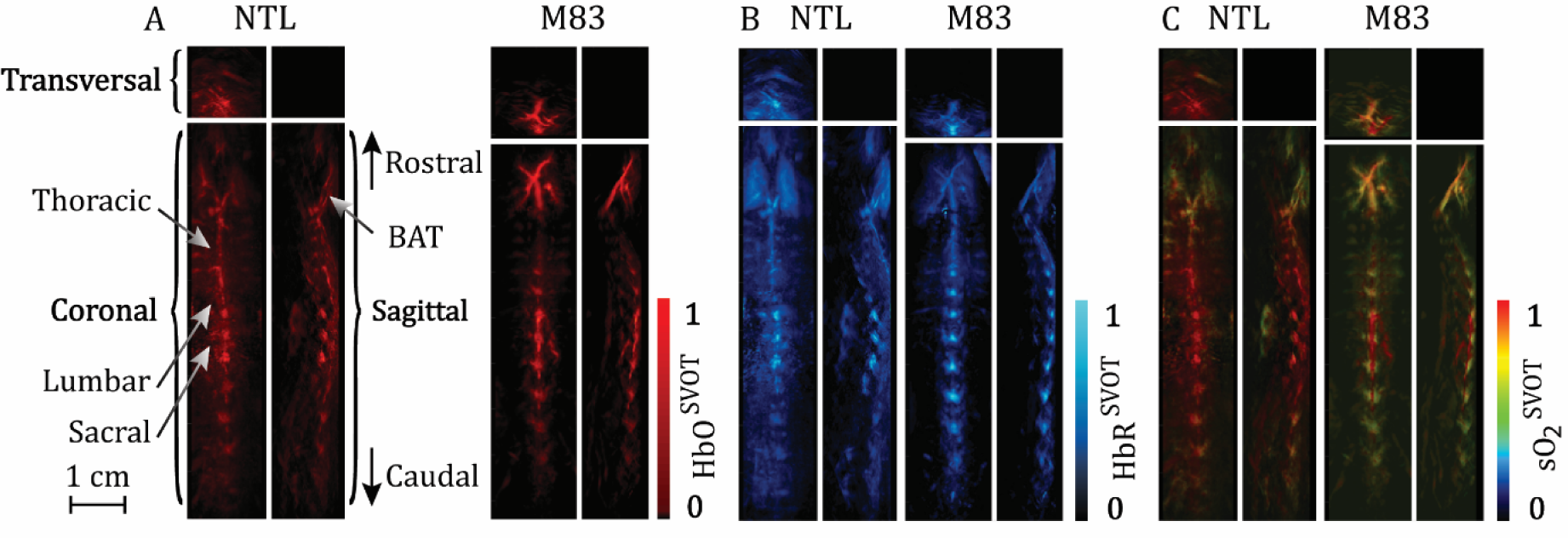
*In vivo* SVOT images of the spinal cords of M83 and NTL (control) mice. A-C) Representative maximum intensity projection images of HbO (A), HbR (B), and sO_2_^SVOT^ (C) in the spinal cords of NTL and M83 mice. Scale bar = 1 cm. BAT, brown adipose tissue.

**Figure 3.**
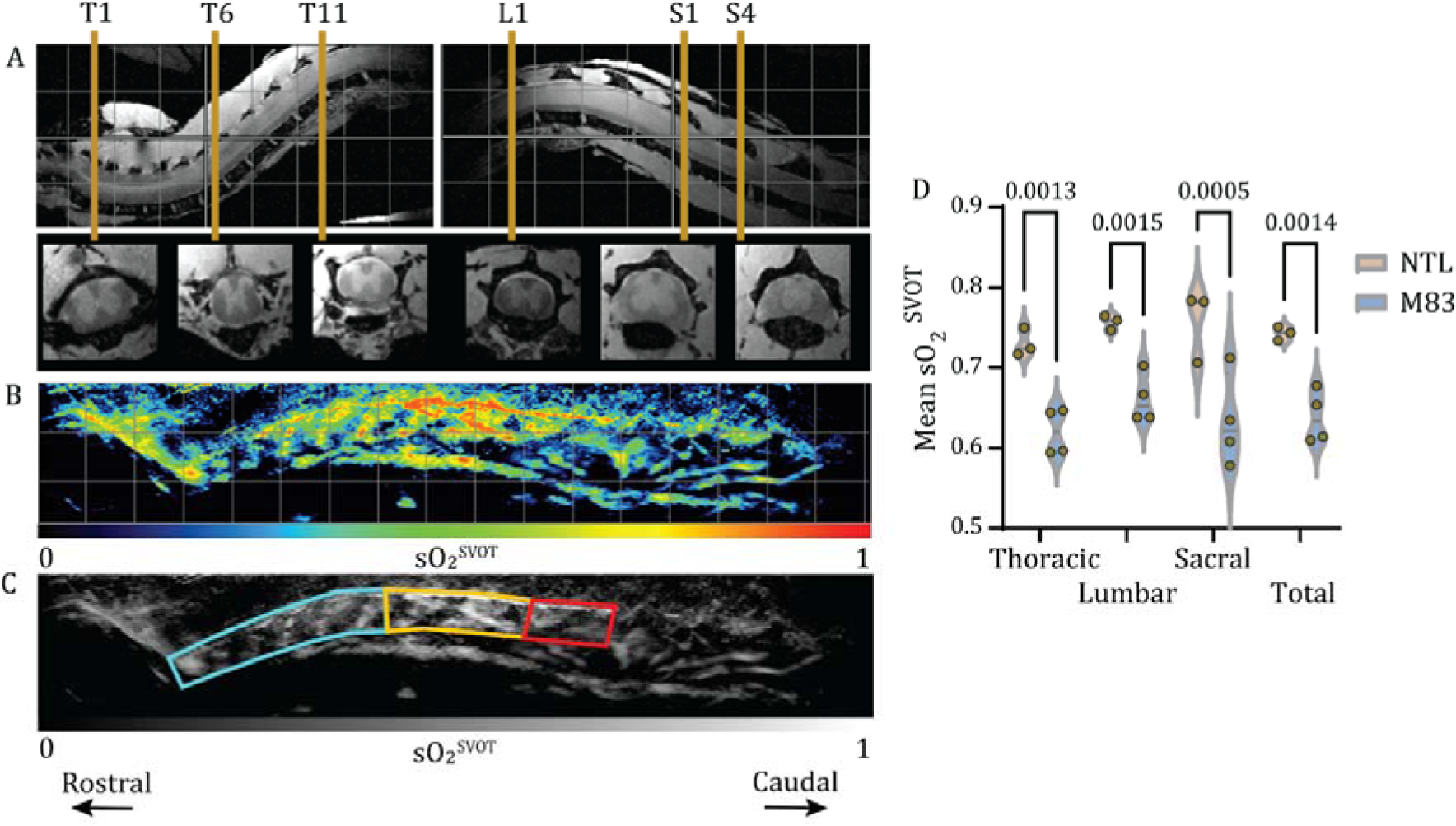
Reduced sO_2_^SVOT^ in the spinal cord of M83 mice compared to NTL mice. A) *Ex vivo* MRI showing sections of the thoracic (T, left), lumbar (L, right) and sacral (S, right) vertebral segments from a M83 mouse. First row: sagittal view, second row: transverse view, grid = 2.5 mm. Transverse views display six landmarks of the spinal cord (T1, T6, T11, L1, S1 and S4) used for segmentation. B) Representative sO_2_^SVOT^ distribution (sagittal projection) from a M83 mouse; grid = 3 mm. C) Representation of the same image transformed into grey scale with regions-of-interest marker for further quantification. Cyan: thoracic, orange: lumbar and red: sacral segments. D) Comparison of mean sO_2_^SVOT^ between M83 and NTL mice in the thoracic, lumbar, and sacral segments and in three combined segments (total). sO_2_ ^SVOT^ scale: 0-1. N = 4 M83 and n = 3 NTLs.

### 2.2 Low oxygenation in the spinal cord is not associated with spinal volumetric reduction

Spinal cord structural organisation was analysed by applying automatic segmentation generated by a deep learning-based model to study potential volumetric changes associated with low spinal oxygenation. To validate the automatic segmentation generated by the deep learning model, we first compared cross-sectional area (CSA) values of grey matter (GM) and white matter (WM) from the model (Fig. 4B) with manually segmented CSAs (Fig. 4A) on the same sections. Spearman’s correlation analysis shown a very strong positive linear relationship between the deep learning and manual segmentation (n = 26, r = 0.9596, p < 0.0001) (Fig. 4C). This indicates that the deep learning-generated GM and WM CSAs model is an efficient and reliable tool to quantify potential structural spinal changes in mice.

**Figure 4.**
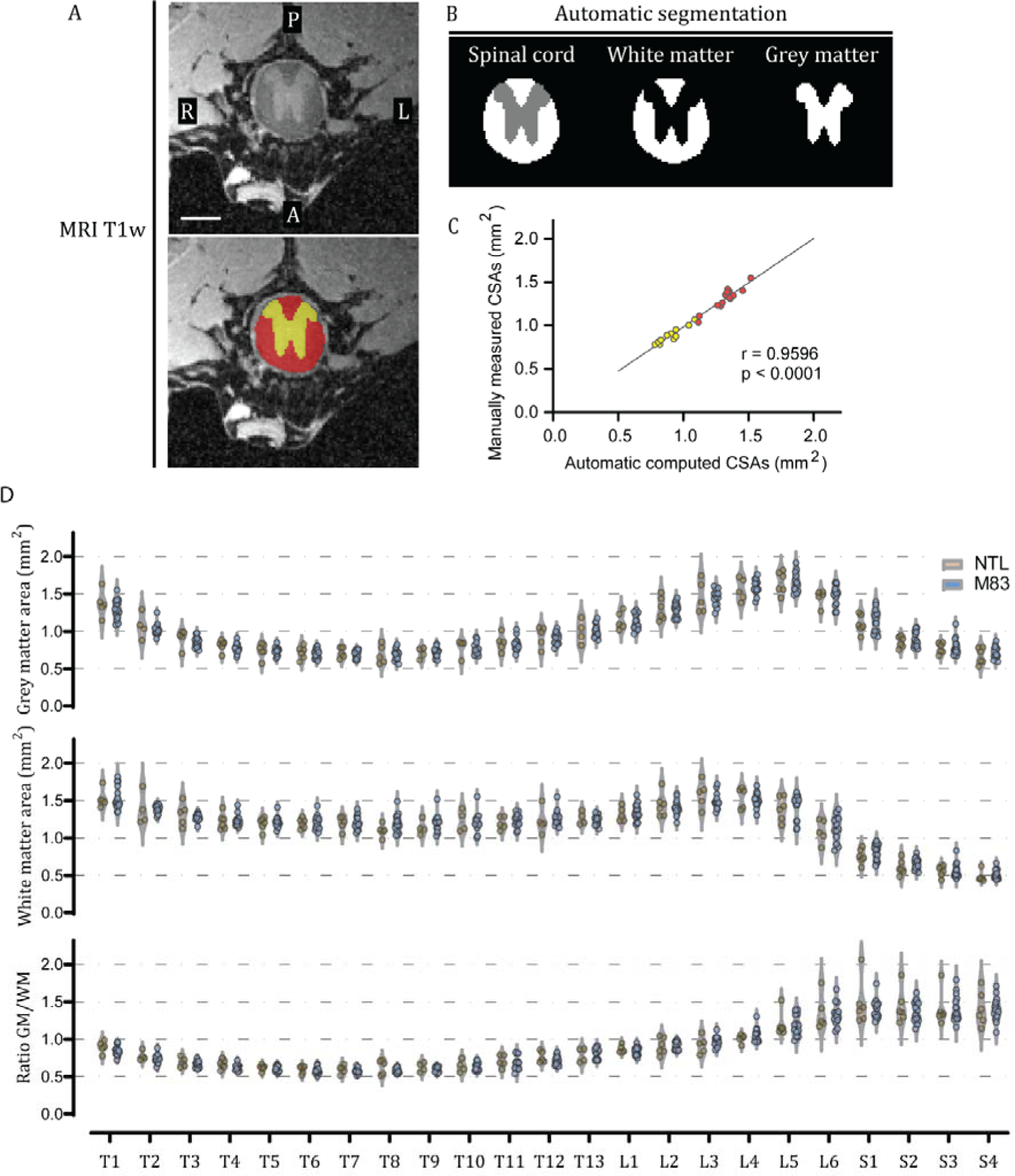
Absence of spinal atrophy in the *ex vivo* T1w MRI images of M83 mice compared to their NTLs (controls). A) Representative transverse section of the thoracic spinal segment. Upper and lower panels show the same image where manual segmentation is depicted in the lower panel. Yellow: grey matter, red: white matter, R: right, L: left, A: anterior, P: posterior; scale bar = 1 cm. B) Representation of deep learning computed segmentation on the same section shown in A. C) Spearman correlation between deep learning-generated and manually segmented CSAs. Yellow and red dots represent values for grey and white matter respectively. D) Comparison of mean grey matter cross-sectional areas (upper panel), white matter cross-sectional areas (middle panel) and grey matter / white matter cross-sectional areas (lower panel) between M83 and NTL mice in the thoracic, lumbar, and sacral segments. N=15 M83 and n=7 NTLs.

Quantification using CSAs indicated that there was no volumetric alteration in the spinal cord GM or WM between M83 mice and their NTLs in any of the thoracic, lumbar, or sacral segments of the spinal cord, as well as in the ratio of GM/WM (Fig. 4D). NeuN staining showed intact Clark’s column in the thoracic and lumbar spinal segments of M83 mice, although the number of motor neurons appear to be relatively low (Supplemental Fig. 1A and B). We further examined whether there was atrophy in the brain of the M83 mice using 9.4T MRI ex vivo. No structural abnormality or volumetric alterations was observed in the M83 mice compared to NTL mice by using MRI (Supplemental Fig. 2A).

Earlier studies have shown a vicious circle between neuroinflammation and hypoxia such as in multiple sclerosis (Yang and Dunn 2019) as well as in neurodegenerative diseases of disrupted brain energy metabolism (Butovsky and Weiner 2018) (van Horssen et al. 2019). Next, we performed staining for astrocytes (GFAP) and microglia (Iba1) in the different spinal segments of M83 mice. No apparent astrocytosis and microgliosis was observed throughout the spinal cord of M83 mice. Both astrocytes and microglia displayed homeostatic non-reactive morphology in the sections examined (Supplemental Fig. 1A and B and Supplemental Fig. 3A and B).

### 2.3 Low oxygenation in the spinal cord is not due to vascular organization impairment

To explore the cause of the reduction in the oxygenation in the spinal cord of M83 mice, we analysed the spinal vascular organization after *in vivo* imaging (Fig. 1C). For this, we first performed immunostaining of the thoracic, lumbar, and sacral regions using the platelet endothelial cell adhesion molecule (CD31), expressed by differentiated endothelial cells (Fig. 5A) (Kecheliev et al. 2023). To assess the vasculature network, we used an ImageJ (Fiji) script to automatically calculate (1) the area fraction of blood vessels, (2) the length of blood vessels and (3) the number of branches and junctions (Rust et al. 2019) based on the CD31 staining. No significant differences were observed in the vasculature area (p=0.1586, p=0.1827 and p=0.1314), in the number of branches (p=0.3300, p=0.3300 and p=0.3127), or in the vascular length (p=0.2262, p=0.1335 and p=0.2001) in the thoracic, lumbar, and sacral segments between M83 and NTL control mice (Fig. 5B, 5C and 5D).

**Figure 5.**
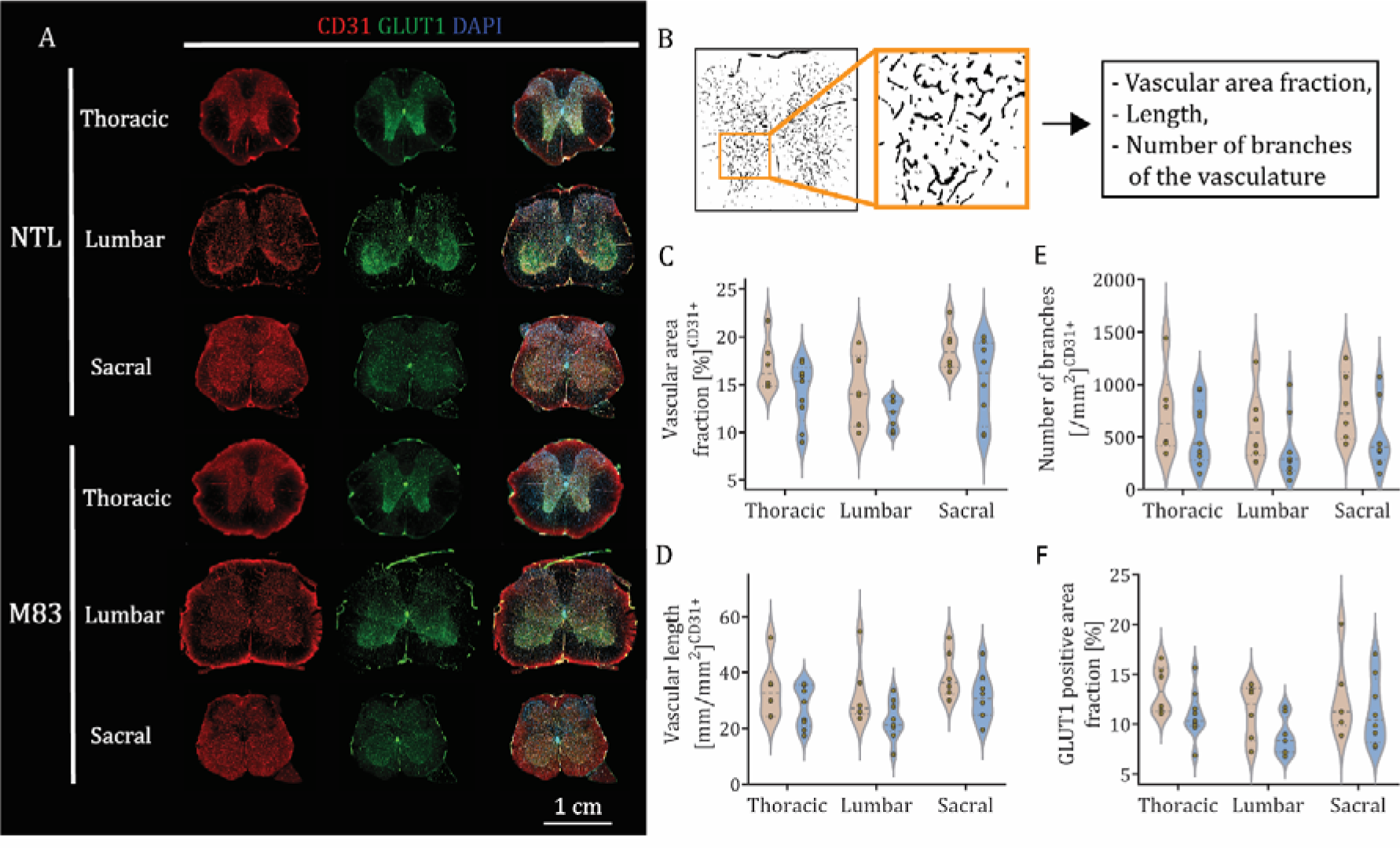
Vascular density, organization and function are not impaired in the spinal cord of M83 mice. A) Representative immunofluorescence images of CD31 (red), GLUT1 (green) and DAPI (blue) at the level of the thoracic, lumbar and sacral vertebras in NTL and M83 mice with a zoom-in showing close colocalization of CD31 and GLUT1; scale bar = 1 cm. B) Representation of the automated vascular organization analysis. C, D and E) Quantitative evaluation of the vascular area fraction, number of branches and blood vessel length in the different spinal segments of both groups. F) Quantitative evaluation of GLUT1-positive area fraction. N=9 M83 and n=6 NTLs.

After analyzing the blood endothelial cell structure, we performed glucose transporter 1 (GLUT1) immunostaining to study the functionality of the vessel in the spinal cord. GLUT1 protein is critical for transporting glucose across the blood-brain barrier to the central nervous system and is highly expressed by endothelial cells (Kong et al. 2024). The GLUT1-positive area fraction was not significantly different between M83 and NTL mice in any of the three spinal segments (p=0.2559, p=0.2559 and p=0.3998, respectively, in the thoracic, lumbar and sacral segments) (Fig. 5E). These results suggest that M83 mice do not exhibit vascular network disruption.

### 2.4 Αlpha-Synucleinopathy throughout the spinal cord in M83 mice

Next, we assessed the distribution of α-syn pathology in the spinal cord sections of M83 mice using anti-pS129 α-syn antibodies. Given the complex finding reported in α-syn staining using different antibodies, we used three widely used antibodies against pS129 α-syn, i.e. EP1536Y clone, 81A clone and MJR-R13 clone. The pS129 α-syn signal (EP1536Y clone) was more abundant at the level of the thoracic, lumbar and sacral spinal segments of the spinal cord in M83 mice compared to NTLs, and mainly in the GM (Fig. 6A). The % pS129-positive area (EP1536Y clone) was significantly higher in M83 mice than NTLs in the thoracic (1.227% vs 0.006%, p=0.001198), lumbar (1.100% vs 0.010%, p=0.001198) and sacral spinal segments (1.192% vs 0.008%, p=0.002525) (Fig. 6B). Next, we calculated the nuclear and the cytoplasmic pS129-positive signals (EP1536Y clone, background subtracted from control immunostaining without primary antibody). The % pS129-positive area (EP1536Y clone) was significantly higher in the nucleus than in the cytoplasmic area in the thoracic (4.08% vs 1.02%, p=0.011673), lumbar (4.21% vs 0.83%, p=0.011673) and sacral spinal segments (3.17% vs 0.96%, p=0.015625) (Fig. 6C). Notably, intranuclear pS129 α-syn staining (EP1536Y clone) was also found in the brain, mainly in the cortex of the M83 mice (Supplemental Fig. 3B). In contrast, using the other two anti-pS129 α-syn antibodies, 81A clone or MJR-R13 clones, no apparent difference in signal intensity and localization of pS129 α-syn was found throughout the spinal cord of M83 mice and NTLs (Supplemental Fig. 4).

**Figure 6.**
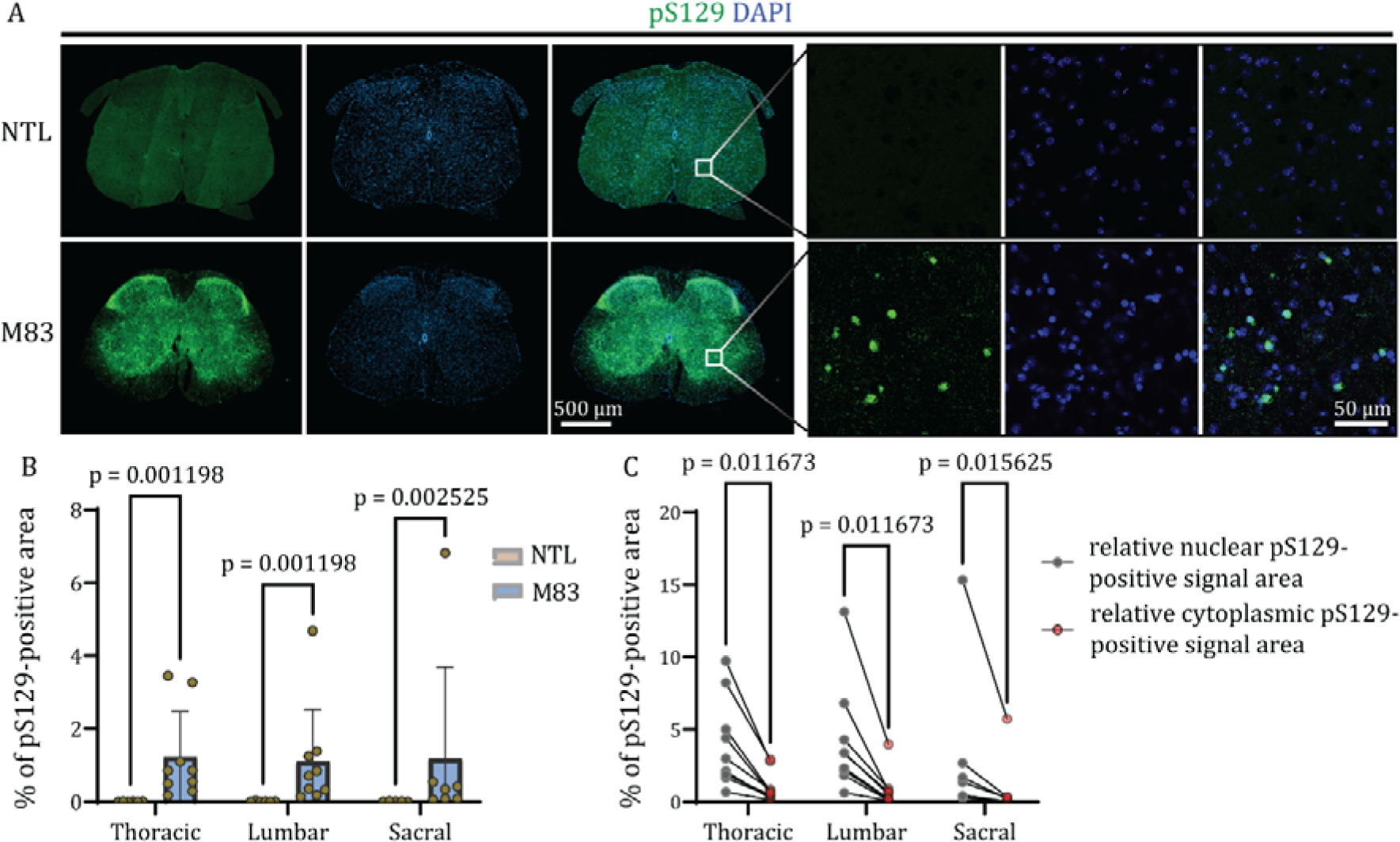
pS129 immunostaining in the spinal cord of M83 and NTL (control) mice using EP1536Y clone. A) Representative immunofluorescence images of colocalised pS129 (green) and DAPI (blue) showing intranuclear pS129-positive signals at the lumbar level of the spinal cord of M83 mice compared to NTL mice. Scale bars = 500 µm and 50 µm. B) Quantitative evaluation of the pS129-positive areas in M83 and NTL control mice, at the level of the thoracic, lumbar, and sacral spinal cord segments. C) Higer levels of relative nuclear pS129-positive areas (grey) than cytoplasmic pS129-positive areas (red dots) in thoracic, lumbar and sacral spinal cord segments. N=9 M83 and n=6 NTLs.

## 3. Discussion

In this work, we demonstrated the preclinical potential of SVOT in measuring the reduced oxygen perfusion in the spinal cord of the M83 mouse model of α-synucleinopathy, with unimpaired GM/WM structure (by MRI) and vascular network in the spinal cord. In addition, we found higher pS129 α-syn accumulation in the spinal cord of M83 mice compared to NTL mice, Higher nuclear than cytoplasmic location of pS129 α-syn was observed in the spinal cord of M83 mice, which might be linked to early pathology and metabolic dysfunction.

Optoacoustic imaging has previously been employed to study brain dynamics (Gottschalk et al. 2019), hemodynamic alterations in neurodegenerative disease (Ni et al. 2018) and stroke (Deán-Ben et al. 2023), as well as other proteinopathies using extrinsic contract agents (Ni et al. 2022; Straumann et al. 2023; Vagenknecht et al. 2022). This method has also been used to detect changes in the spinal cord of an ischemic stroke mouse model (Ni et al. 2023), as well as in an experimental autoimmune encephalomyelitis (EAE) model of MS (Ramos-Vega et al. 2022). To our knowledge, the current study is the first optoacoustic imaging study in PD mouse models. We demonstrated mapping of oxygenation abnormalities in the spinal cord of PD mice with sub 100-micron spatial resolution. SVOT is a state-of-the-art optoacoustic tomography imaging system for preclinical volumetric 3D imaging of the whole-body of small animals with scalable spatio-temporal resolution (Kalva et al. 2023b). While other cross-sectional optoacoustic imaging systems can potentially be utilized for whole-body imaging via scanning of a circular ring-shaped array transducer (Kalva et al. 2023a, b), the proposed SVOT configuration attains true volumetric (3D) imaging capacity on a whole-body scale with nearly isotropic resolution, making it highly suitable for accurate signal quantification in the spinal cord.

By using *in vivo* SVOT, we showed a 13-16% reduction in the oxygenation level in the thoracic, lumbar, and sacral spinal segments of asymptomatic M83 mice compared to NTLs. Lower spinal cord [^18^F]fluorodeoxyglucose hypometabolism has been observed in M83 mice at 9 months of age (Mondal et al. 2021). The low oxygen saturation found in the spinal cord of M83 mice is in line with previous studies on cerebral α-syn-associated mitochondrial degeneration in this model when treated with the pesticide paraquat (which is absent in the wild-type human α-syn M7 line) (Norris et al. 2007). In the M83 mouse model, the administration of the iron chelator clioquinol compound that stabilized functional hypoxia-inducible factor-1α, led to improvement in motor function (Finkelstein et al. 2016). This indicates an important role of hypoxia in the motor function of PD rodent model.

Hypoperfusion is hypothesized to contribute to neurodegeneration by lowering the neuronal energetic supply in human (Hernandez-Gerez et al. 2019) (Wolters et al. 2017). In mouse models, chronic cerebral hypoperfusion led to WM lesions in the brain (Shibata et al. 2004). In aditional, neuronal functional deficits are associated with spinal cord hypoxia in an EAE model. To study potential structural changes and volumetric atrophy associated with oxygenation impairments, we developed a deep learning model that able to use 3D MRI images as input and automatically segment the GM and the WM. Upon validation of the model, no atrophy was found in the thoracic, lumbar or sacral spinal segments of M83 mice compared to their NTLs. No difference was observed tin the GM, WM areas, or WM/GM ratio (computed with CSAs) for each segment bewteen M83 and NTL mice. An earlier study has shown spinal axonal degeneration with deteriorating myelin in 12 month-old M83 mice (Giasson et al. 2002). Previous studies have shown impaired neurons in the spinal cord of rodent models of 1-methyl-4-phenyl-1,2,3,6-tetrahydropyridine or rotenone-induced experimental parkinsonism by immunostaining (Chera et al. 2002; Chera et al. 2004; Samantaray et al. 2007). Here we did not observe apparent alterations in the NeuN staining of the spinal cord slices of M83 mice. One possibility is that asymptomatic M83 mice in this study had oxygen saturation impairments before presenting strong markers of neurodegeneration. *Ex vivo* MRI might not be sensitive enough to detect axonal degeneration at this stage.

The histopathology in the brain, and behavior abnormality development of the transgenic M83 PD mouse model have been well characterized (Giasson et al. 2002). Although the developed motor symptoms are thought to be an outcome of pyramidal or motor neurodegeneration rather than the result of nigrostriatal degeneration in this model (Benskey et al. 2016), little is known about the related pathological processes in the spinal cord. Very few in vivo or *ex vivo* MRI or optical studies have been performed in the spinal cord of PD animal models (Chu et al. 2020; Ni 2022). Functional and structural MRI has been performed on the spinal cord of wild-type (Bilgen et al. 2005; Saito et al. 2012), tau mice (Sartoretti et al. 2022), amyotrophic lateral sclerosis mice (Gao et al. 2020; Gatto et al. 2018) and experimental autoimmune encephalomyelitis model. To our knowledge, our study is the first MRI study investigating the alterations in spinal cord of PD animal model.

To understand the possible reasons for the observed reduction in sO_2_ ^SVOT^ in M83 compared to NTLs, we investigated possible causes, including alterations in the vasculature, α-syn level as well as glial activation. Our *ex vivo* analysis of spinal vascular organization and functionality suggested that the vascular network was not impaired. Quantification of vascular density and organization with CD31 expression did not show any significant differences in the vascular area fraction, number of branches or blood vessel length between M83 and NTL mice. GLUT1 expression did not display significant changes between the two strains, suggesting that vessel functionality was also not impaired and was not the cause of the reduction in oxygen saturation. It is also important to note that GLUT1 expression can be enhanced under hypoxic conditions, mainly in tumor environment, which was not the case in our study, in which the impairment in oxygenation is probably only mild (Bukkuri et al. 2022).

Neuroinflammation is closely linked to cerebral energy metabolism impariments in neurodegenerative diseases. Recently, induction of neuroinflammation in a mouse model of Alzheimer’s disease has been shown to elicit reductions in cerebral intravascular oxygen and increases in oxygen extraction in the brain (Davies et al. 2013; Liu et al. 2024). The neuroinflammatory pathology in the EAE mouse model leads to hypoxia accompanied by reductions in spinal vascular perfusion (Ramos-Vega et al. 2022). In this study, we did not observe any sign of inflammation in the spinal cord of M83 mice by using immunohistochemistry, as astrocytes and microglia do not display reactive phenotypes. This is in line with earlier study showing gliosis only in the symptomatic M83 and M83 mice injected with α-syn preformed fibrils (Sorrentino et al. 2018). Therefore, further study is needed to elucidate the cause of low oxygen saturation in the spinal cord of M83 mice.

pS129 α-syn has been the focus in understanding α-synucleinopathies because it is one of the most robust pathological markers of early α-syn aggregation, with almost all the aggregates containing this posttranslational modification (Anderson et al. 2006; Fujiwara et al. 2002). On the other hand, pS129 α-syn is known to play a physiological role at the synaptic level in healthy brains and it triggers protein-protein interactions(Parra-Rivas et al. 2023; Ramalingam et al. 2023). Accumulation of pS129 α-syn has also been observed in the context of oxygen intake and utilization disorders in cells, as well as in middle cerebral artery occlusion rodent models,(Kim et al. 2016; Unal-Cevik et al. 2011) and in patients with obstructive sleep apnea syndrome (Chen et al. 2014). Nuclear translocation of pS129 α-syn has also been shown following transient focal ischemia in a mouse model of stroke (Kim et al. 2016). Our study revealed an increase in the pS129 α-syn-positive signal within the thoracic, lumbar, and sacral spinal segments in M83 mice compared to their NTLs. Immunofluorescence staining revealed that the main location of pS129 α-syn was in the vicinity of the nucleus. Several studies have suggested a possible nuclear role of α-syn in modulating gene expression, possibly by binding to DNA (Miller et al. 2007), in mouse and cellular models (Kontopoulos et al. 2006; Outeiro et al. 2008; Siddiqui et al. 2012) as well as in human PD brains (Garcia-Esparcia et al. 2015; Siddiqui et al. 2012). Studies have shown the presence of nuclear and perinuclear pS129-α-syn-positive signals on brain slices from transgenic ((Thy1)-[A30P]αSYN) or α-syn fibril-injected rodent models using high-affinity antibodies against the specific α-syn phosphorylated at the Ser129 site (EP1536Y or D1R1R)(Schell et al. 2009) (Delic et al. 2018; Elfarrash et al. 2021). pS129 α-syn has been shown to modulate the nuclear localization of the wild-type form of α-syn, affecting gene expression, suggesting an important role of the phosphorylation status as well as the different α-syn species within the nucleus (Pinho et al. 2019).

The presence of pS129 α-syn in the nucleus remains controversial as numerous antibodies targeting this posttranslational modification of α-syn have been developed with off-target, as well as non-specific binding and disparity in staining signal using these antibodies as reported earlier (Delic et al. 2018; Huang et al. 2011; Kumar et al. 2020; Lashuel et al. 2022). It is noted that α-syn staining appeared to be antibody (clone-dependent) in the spinal cord of M83 mice, with negative staining observed using two other clones targeting pS129 (81A and MJR-R13). Nevertheless, the clone EP1536Y is one of the most robust and widely used antibodies to study alpha-synuclein pathology (Delic et al. 2018). The difference in the pS129-positive signal observed and quantified between M83 mice and NTLs was genotype dependent in our study. These observed alterations can be very useful for studying early pathological changes in this model of PD. Further mechanistic studies on the potential association between low oxygen saturation and intranuclear pS129 α-syn, and the consequences of this nuclear location of pS129 α-syn will be informative.

Heterozygous and homozygous M83 mice develop motor impairment at 22-28 and 8-16 months of age respectively. We focused on studying early changes in asymptomatic mice of both hetero- and homozygous M83 mice for the expression of the human mutated A53T α-syn. Similar results in the spinal perfusion, spinal volume and pS129 load was observed in M83 hetero- and homozygous mice.

There are several limitations in the present study: 1) unmatched *in vivo* and *ex vivo* sample sizes and, sex imbalance in the samples (mice with pigments on the skin inside the region of interest after shaving were excluded from the *in vivo* study and were only used for *ex vivo* analysis); 2) spectral colouring effect due to wavelength dependent light attenuation (Cox et al. 2012; Hochuli et al. 2019), which can potentially be reduced by correcting for light attenuation with depth (Chen et al. 2022); 3) the cervical part of the spine region displayed very little or no signal and thus could not be measured due to the high absorption of the BAT. Further studies are needed to elucidate the cause underlying the reduction in sO_2_ in the spinal cord of M83 mice. 4) We did not further segment GM and WM in the analysis of immunostaining results.

## 4. Conclusion

In conclusion, we demonstrated the use of a non-invasive high-resolution imaging tool to study oxygenation *in vivo* in the spinal cord of the PD M83 mouse model. SVOT has the potential to measure *in vivo* spinal cord sO_2_, which is an early pathological marker preceding the motor deficits in M83 mice. *Ex vivo* high-resolution MRI combined with deep learning automated quantification has further shown that low spinal cord oxygen saturation was not associated with volumetric atrophy in this model of α-synucleinopathy. This new approach thus might facilitate our understanding of metabolic and potential structural changes in the spinal cord of rodent models of PD and the monitoring of treatment effects.

## 5. Methods

### Animal model

In total, 22 M83 mice (14 homozygous and 8 heterozygous) between 9 and 12 months of age overexpressing human mutated A53T α-syn under the mouse prion promoter (C57Bl/C3H background) (Giasson et al. 2002; Straumann et al. 2023) and 13 age-matched non-transgenic littermates (NTLs) of both sexes (M83 10/12 males/females and NTLs 6/7 males/females) were used for different purposes throughout the study. Four M83 (2/2 males/females) and three NTLs (0/3 males/females) were scanned using the *in vivo* SVOT system, 15 M83 (9/6) and 7 NTLs (2/5) were imaged *ex vivo* with MRI, and 9 M83 (3/6 males/females) and 6 NTLs (4/2 males/females) were included in the histological tissue analysis. None of the animals displayed abnormal posture, seizures, or paralysis, as can be observed in end-stage aged M83 mice. All the M83 mice were asymptomatic. Animals were housed in individually ventilated cages inside a temperature-controlled room under a 12-h dark/light cycle. Pelleted food (3437PXL15, Cargill) and water were provided ad libitum. All the experiments were performed in accordance with the Swiss Federal Act on Animal Protection and were approved by the Cantonal Veterinary Office Zurich (ZH024/2021).

### Spiral volumetric optoacoustic tomography (SVOT)

The SVOT system employs a Nd:YAG-pumped optical parametric oscillator laser (SpitLight, Innolas Laser GmbH, Krailling, Germany) as an excitation light source. It delivers < 10 ns duration pulses at a repetition rate of 10 Hz over a broad tunable wavelength range (680–1250 nm) with per-pulse energies up to ∼180 mJ. Five wavelengths (700, 730, 760, 800 and 850 nm) were employed repeatedly in succession (20 averages per wavelength) to efficiently unmix the HbO, HbR and total hemoglobin components using the spectral absorption profiles of hemoglobin in oxygenated and deoxygenated forms (Jacques 2013). The light beam was guided through a custom-made fiber bundle (CeramOptec GmBH, Bonn, Germany) placed at a radial distance of 40 mm from the center of a custom-made spherical array. The output bundle created a Gaussian illumination profile with a size of 10 mm at full width at half maximum on the mouse skin surface. The optical fluence on the skin surface was maintained within safe limits according to the American National Standards Institute regulations throughout all experiments (American National Standards and Laser Institute of 2022). A custom-made spherical array, comprising 512 distinct piezoelectric sensor elements, each with a surface area of ∼7 mm², a central detection frequency of 7 MHz, and a detection bandwidth of ∼85% (spanning from 2.6 to 8.6 MHz at the full width at half maximum), was employed to collect the optoacoustic responses (Kalva et al. 2023a). These elements were organized on a hemispherical surface with a radius of 40 mm and an angular coverage of 110 degrees (0.85π solid angle). Simultaneously, the optoacoustic signals were digitized at a rate of 40 Mega samples per second using a tailored parallel data acquisition unit, Falkenstein Mikrosysteme GmBH, Taufkirchen, Germany). This data acquisition unit was synchronized with the laser’s Q-switch output and connected to a PC via a 1 Gb/s Ethernet link for data storage and subsequent analysis. The data acquisition process was performed under a custom MATLAB interface (Version R2020b, MathWorks Inc., Natick, MA, USA) running on a PC. Mice (n=4 M83 and n=3 NTLs) were anesthetized with isoflurane (induction: 4% and maintenance: 1.5-2%) and imaged following protocol previously described (Kalva et al. 2023b) and maintained under constant anaesthesia and at 36 ± 0.5□°C throughout the experiments.

### SVOT image reconstruction and spectral unmixing

At each scanning position of the spherical array on the back of the mouse, signals were initially averaged 20 times for each wavelength (700, 730, 760, 800 and 850 nm), band-pass filtered between 0.1-12 MHz, and eventually deconvolved with the impulse response of the spherical array sensing elements. Next, a GPU-implemented back-projection (BP) reconstruction technique was employed for the reconstruction of individual volumetric frames, with each US sensing element split into 16 sub-elements (Kalva et al. 2021). Whole-spinal cord images were obtained by stitching individual reconstructed volumes at each corresponding position of the spherical array. To quantify the HbO- and HbR levels in the spinal cord, a linear spectral unmixing algorithm was employed (Deán-Ben et al. 2023; Gottschalk et al. 2019). Before unmixing, the reconstructed whole-spinal cord volumes for each wavelength were normalized to the respective optical fluence values. The oxygenation values rendered with SVOT on a voxel-by-voxel basis (sO_2_^SVOT^) were then calculated as (HbO/(HbO + HbR))*100. All the image reconstruction and processing steps were performed in MATLAB (MathWorks, Natick, MA, USA). maximum intensity projection images (sagittal view) of the grey scale sO_2_^SVOT^ from the MATLAB files were captured and used for analysis. Regions-of-interest (ROIs) were drawn on these images using ImageJ (NIH) to measure the mean sO_2_ ^SVOT^ signal intensities in the spinal cord segments. using the atlas of the mouse spinal cord (Watson et al. 2009) as reference.

### Sample Preparation

Immediately following SVOT imaging, M83 and NTLs were intracardially perfused under deep anaesthesia with 0.1 M phosphate buffered saline (PBS, pH 7.4) followed by 4% paraformaldehyde in 0.1 M PBS (pH 7.4). Mouse head and vertebral column samples were postfixed in 4% paraformaldehyde in 0.1 M PBS for 6 days and stored in 0.1 M PBS (pH 7.4) at 4 °C as described earlier (Sartoretti et al. 2022).

### Ex vivo MRI of the M83 mouse head and spinal cord

Mouse head and vertebral column samples were placed in a 15 ml centrifuge tube filled with perfluoropolyether (Fomblin Y, LVAC 16/6, average molecular weight 2,700; Sigma□Aldrich, USA) (Massalimova et al. 2021). Data were acquired on a BioSpec 94/30 preclinical MRI scanner (Bruker BioSpin AG, Fällanden, Switzerland) with a cryogenic 2×2 radio frequency phased-array surface coil (overall coil size of 20×27 mm^2^). The coil system operated at 30□K for reception in combination with a circularly polarized 86□mm volume resonator for transmission (Ni et al. 2020). For the spinal cord, a structural T1w scan was acquired with a 3D multishot echo planar imaging sequence (4 shots) with a field of view =□25 mm × 10 mm × 10 mm and matrix dimension =□500 × 200 × 200, resulting in a nominal voxel resolution of□50 × 50 × 50 μm. The following imaging parameters were chosen: echo time = 8 ms, repetition time =□50 ms, number of averages□=□4 The total acquisition time was 2 h 28 min for each segment (Sartoretti et al. 2022).

For the brain, a structural T1 scan was acquired with a 3D multishot echo planar imaging sequence (4 shots) with a FOV of 18 × 12 × 9 mm and matrix dimension of 180 × 120 × 90, resulting in a nominal voxel resolution of 100 × 100 × 100 μm. Structural T2 scans were acquired using a T2-Turbo rapid acquisition with relaxation-3D sequence with a field of view of 15 × 15 × 8 mm and matrix dimension of 150 ×150×80, resulting in a nominal voxel resolution of 100 × 100 × 100 μm, echo time □=□ 49 ms, repetition time =□ 2000 ms, number of averages□=□1, acquisition scan time □= 52 min 48 sec (Kecheliev et al. 2023).

### MRI data postprocessing and analysis

ITK-SNAP software (v4.0.1, Penn Image Computing and Science Laboratory - PICSL, University of Pennsylvania, Philadelphia, PA) was used to inspect and manually reorient the MR images of the spinal cord. By using an anatomical atlas (Watson et al. 2009) for guidance, representative axial T1w image sections matching the 23 spinal segments from the thoracic, lumbar and sacral parts (T1-T13, L1-L6 and S1-S4) were identified. GM and WM in the spinal cords were automatically segmented using a deep learning-based model trained on manually annotated slices and improved by active learning (https://github.com/ivadomed/model_seg_mouse-sc_wm-gm_t1). The model is based on a 3D nnUNet architecture (Isensee et al. 2021), which takes as input a 3D MRI image and outputs a 3D segmentation file with values 1 for GM, 2 for WM and 0 for background. The GM and WM CSAs were then extracted using the sct_process_segmentation command from the Spinal Cord Toolbox (De Leener et al. 2017) (v6.1) for all 23 identified segments. The average of five CSAs from 5 consecutive sections was calculated per segment and per mouse. GM and WM CSAs as well as WM/GM ratios were further analysed to study potential spinal volumetric atrophy.

Validation of the deep learning method was performed by comparing the automatically computed CSAs (without angle correction) with the manually segmented CSAs from the same 13 thoracic sections (T1-T13) for one mouse. GM and WM CSAs were analysed to study potential volumetric alterations.

### Immunofluorescence and immunohistochemical staining

Mouse brain and spinal cords were extracted and placed in 0.1 M PBS for 24 h in 15% sucrose and in 30% sucrose in 0.1 M PBS until the tissue sank or for a maximum of 5 days. Coronal brain sections (30 µm) were obtained using a HM 450 Epredia sliding microtome (Thermo Scientific, UK). Spinal cords were embedded in OCT in SpineRacks (Fiederling et al. 2021) made of natural PVA and printed with the Ultimaker S5 3D printer (Ultimaker B.V.; Geldermalsen, Netherlands). Transverse sectioning (30 µm) was performed using a Leica CM1900 cryostat (Leica Biosystems, Germany). After cutting, free floating sections were stored in 0.1 M PBS + 0.1% sodium azide at 4 °C.

For immunofluorescence staining, sections were rinsed three times in 0.1 M PBS for 10 min, followed by a 1 h incubation in 0.1 M PBS, 5% v/v normal donkey serum (NDS), 5% v/v normal goat serum (NGS), and 0.5% v/v Triton-X for blocking and permeabilization. Primary antibodies against NeuN, GFAP, CD31, GLUT-1 and pS129 (EP1536Y, 81A and MJR-R13 clones), were incubated overnight at 4°C (in 0.1 M PBS, 3% v/v NDS, 3% v/v NGS and 0.3% v/v Triton-X) (Kecheliev et al. 2023; Sobek et al. 2023). The sections were then rinsed three times for 10 min each in 0.1 M PBS before being incubated for 2 h in species specific secondary antibodies (suspended in 0.1 M PBS with 3% v/v NDS, % v/v NGS and 0.3% v/v Triton-X). The tissues were then counterstained with 4’,6-diamidino-2-phenylindole (DAPI) and mounted on microscope slides in Prolong Diamond antifade mounting medium.

For immunochemistry staining, sections were rinsed three times in 0.1 M PBS for 10 min, followed by a 30 min incubation in 0.1 M PBS, 1% H_2_O_2_, before another three times washing and 1 h incubation in 5% v/v NGS, 0.5% v/v Triton-X for blocking and permeabilization. Primary antibodies against Iba1 suspended in 0.1 M PBS, 3% v/v NGS, 0.3% v/v Triton-X, was incubated over night at 4°C. The sections were then rinsed three times for 10 min each in 0.1 M PBS before being incubated in biotinylated anti-mouse secondary antibody (suspended in 0.1 M PBS with 0.5% v/v Triton-X) for 2 h. The sections were then incubated in avidin-biotin complex– horseradish peroxidase for 1 h at room temperature. The sections were developed with 0.025 % 3,3′-diaminobenzidine and 0.05 % H_2_O_2_ in triphosphate buffered saline (TBS, pH 7.4) for 3 min. After being mounted on slides, the sections were dehydrated by ascending alcohol series 70%, 90%, and 100% (twice each) and Roticlear for 2 min. Coverslip was finally mounted with roticlear/rotimount mounting medium.

### Image acquisition and microscopic analysis

For vascular organization analysis, platelet endothelial cell adhesion molecule, also known as CD31 staining and GLUT1 staining of two or three sections per segment (thoracic, lumbar and sacral) were imaged at 10× magnification using a TCS SP8 confocal laser scanning microscope (Leica, Germany). For CD31 staining, the vascular area fraction, length, and number of branches of the spinal cord section were assessed by using automated analysis as previously described (Rust et al. 2019). For GLUT1 staining, the area fraction of the spinal cord section was analysed. Three or four pictures per segment (thoracic, lumbar and sacral) were taken at 63× magnification using a confocal laser scanning microscope to study the pS129-positive signal in the of the spinal cord section. Automatic quantification of the colocalization of pS129-positive pixel with DAPI-positive pixel was performed with ImageJ (Fiji, NIH) to quantify its presence in the vicinity of the nucleus. To study the relative nuclear fraction of pS129 α-syn, we divided the pS129-positive signal colocalized with DAPI by the DAPI-positive area. This fraction was named “relative nuclear pS129-positive signal area”. On the other hand, the remaining fraction of the pS129-positive signal was divided by the total area of the image subtracted by the DAPI area to analyse the fraction occupied by this signal in the cytoplasmic compartment (“relative cytoplasmic pS129-positive signal area”). Representative images of CD31, GLUT1 and pS129 staining of the spinal cord section were taken using Zeiss Axio Scan Z1 slidescanner (Carl Zeiss Meditec AG, Germany) at 20× magnification and TCS SP8 confocal laser scanning microscope at 63× magnification. Representative images of GFAP and NeuN staining were taken using TCS SP8 confocal laser scanning microscope at 10× and 20× magnification. Representative images of Iba1 staining were taken using Zeiss Axio Scan Z1 slidescanner at 20× magnification.

### Statistical analysis

All statistical analyses were performed using GraphPad Prism 9.5.0 (GraphPad Software, Inc., USA). Data distributions were first tested for normality by visually assessing the histograms and Shapiro-Wilk test. For distributed data (Fig. 3 and 5), groups were compared using two-way analysis of variance (ANOVA) followed by Holm-Sidak’s multiple comparison post hoc analysis. The data are presented as violin plots with all the points, quartiles and median. data were not normally distributed. Unpaired Mann-Whitney test and Wilcoxon matched-pairs signed rank test, followed by Holm-Sidak’s multiple comparison *post hoc* analyses have been performed for Fig. 6B and 6C as data was not normally distributed. Violin plots display the mean and bar plots present the mean ± standard deviation. Significance was set at ∗p < 0.05.

## Declaration

### Data and code availability

All raw data are available upon request to corresponding authors. Code for deep learning-based MRI analysis is available at https://github.com/ivadomed/model_seg_mouse-sc_wm-gm_t1.

## Author contributions

BC, SK, DR and RN conceived and designed the study. BC, SK and DR developed the SVOT system. SK and XLDB performed the SVOT experiments. AT and LMH performed the MRI experiments. PLB, JN and JCA developed the deep learning-based model for spinal cord segmentation. BC, PLB, JN and JCA contributed to the extraction of relevant data from automatic segmentation. ID performed genotyping. BC performed histopathological immunostaining. BC, SK, MK, RW, PG and R.N. analysed the data. BC wrote the first draft. All authors contributed to the revision of the manuscript. All authors read and approved the final manuscript.

## Acknowledgements

The authors would like to thank Center for Microscopy and Image Analysis (ZMB), University of Zurich; Michael Reiss, Mark-Aurel Augath and Diana Kindler at Institute for Biomedical Engineering, ETH Zurich and Daniel Schuppli, and Uwe Konietzko at Institute for Regenerative Medicine, University of Zurich. for technical assistance.

---

**Suppl Fig 1.**
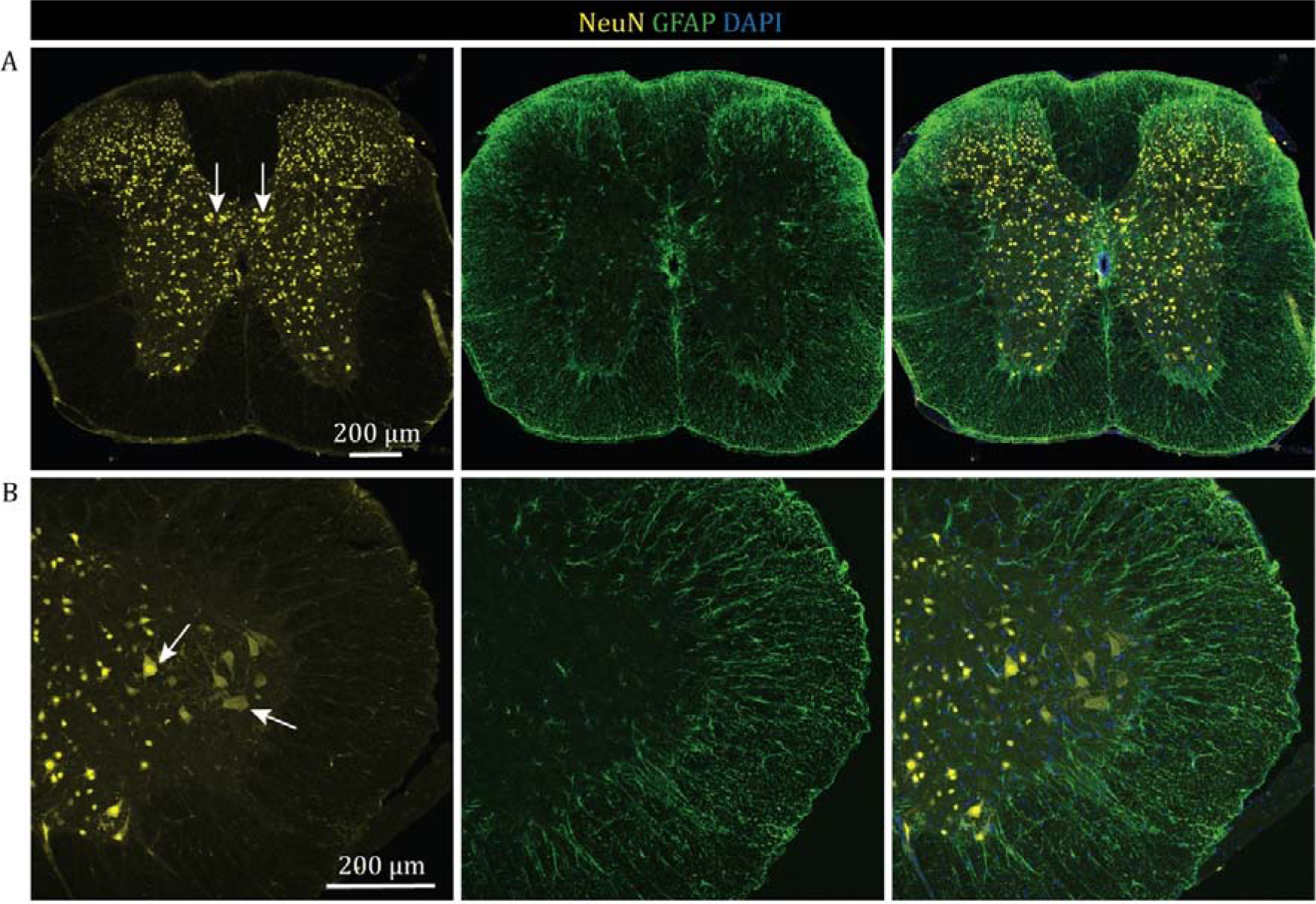
Neuronal and astroglial staining in the spinal cord of the M83 mouse model. A) Representative immunofluorescence images of NeuN (yellow) and GFAP (green) in the thoracic spinal region of M83 mice showing intact Clarke’s column (white arrows) and GFAP-positive astrocytes. B) Representative immunofluorescence images of NeuN and GFAP in the lumbar spinal region of M83 mice showing intact motor neurons (white arrows) and GFAP-positive astrocytes. Nuclei were counterstained using DAPI (blue). Scale bars = 200 µm.

**Suppl Fig 2.**
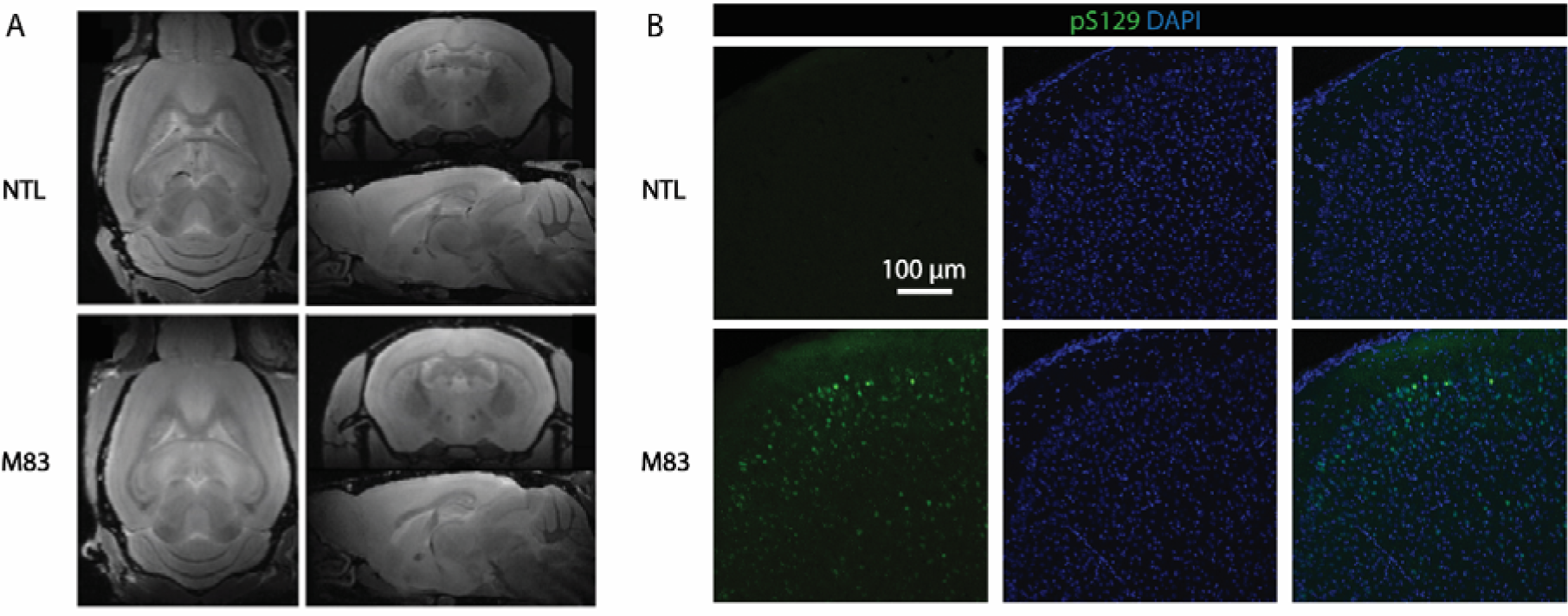
No regional atrophy in the brains of M83 mice compared to NTL mice, despite presence of cerebral pS129-positive (clone EP1536Y) α-Synuclein accumulation. A) High-resolution *ex vivo* T1w MR image acquired at 9.4 T. No apparent atrophy was observed in the brains of M83 mice compared to NTL mice. B) Representative immunofluorescence images of colocalised pS129 (green) and DAPI (blue) showing intranuclear pS129-positive signals in the cortex of the M83 mouse brain, which is absent in the NTL mouse brain. Scale bar = 100 µm.

**Suppl Fig 3.**
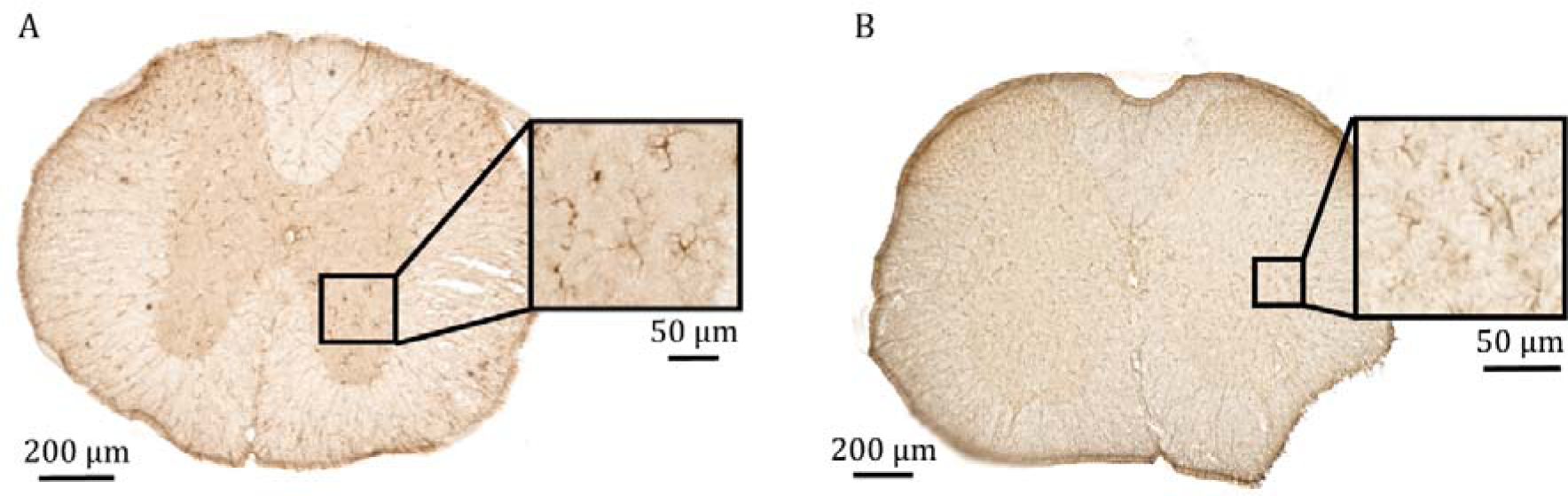
No apparent microglia or astrocyte activation in the spinal cord of M83 mice. A) Representative immunohistochemistry images of Iba1-positive microglia in the thoracic spinal segment of the M83 mouse model. B) Representative immunohistochemistry images of GFAP-positive astrocytes in the thoracic spinal segment of the M83 mouse model. Scale bar = 200 µm and 50 µm.

**Suppl Fig 4.**
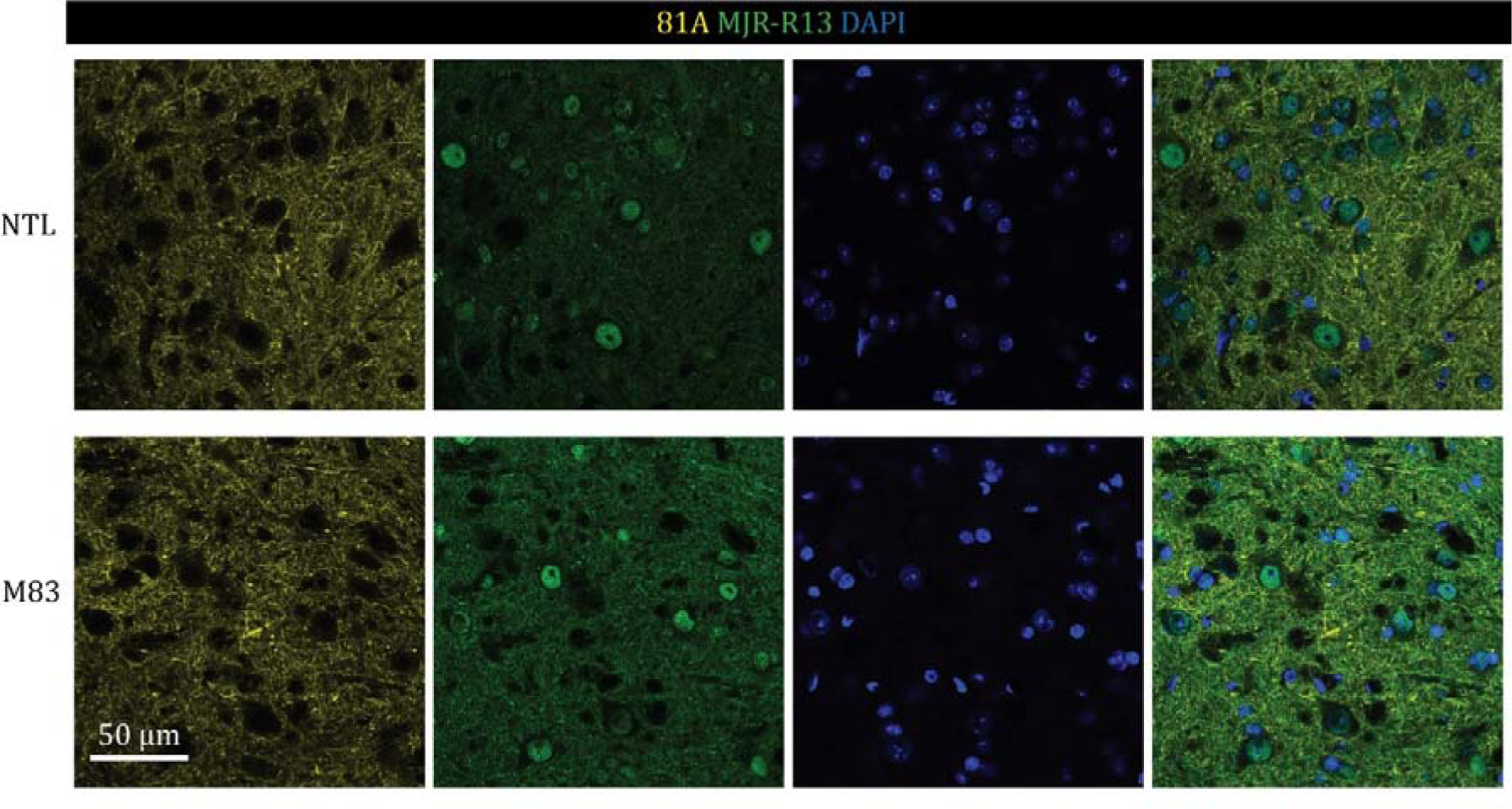
No apparent difference in pS129 expression in the spinal cord of M83 mice compared to NTL mice revealed by the antibodies 81A and MJR-R13 clones. Representative immunofluorescence images of pS129 targeted with 81A (yellow) and MJR-R13 (green) clones showing similar signal in the thoracic spinal cord segment of the M83 mouse model and NTLs. Nuclei were counterstained using DAPI (blue). Scale bar = 50 µm.

**Supplementary Table 1.** Antibodies, chemicals, and materials used.

| Reagent | Dilution | Source | Identifier |
| --- | --- | --- | --- |
| OCT Mounting media |  | VWR chemicals | 361603E |
| NDS |  | Interchim | UP77719A K |
| NGS |  | Jackson ImmunoResearch | 005-000-121 |
| Triton X-100 |  | Merck | RES3103T-A101X |
| Anti-CD31, MEC 13.3 (rat) | 1/50 | BD Pharmingen | 550274 |
| Anti-GLUT-1, 5B12.3 (mouse) | 1/1000 | Merck Millipore | MABS132 |
| Anti-Iba1 (rabbit) | 1/500 | Dako | 019-19741 |
| Anti-alpha-synuclein pS129, 81A (mouse) | 1/1000 | Merck Millipore | MABN826 |
| Anti-alpha-synuclein pS129, MJR-R13 (rabbit) | 1/1000 | Abcam | ab168381 |
| Anti-NeuN, A60 (mouse) | 1/100 | Millipore | MAB377 |
| Anti-GFAP (rabbit) | 1/200 | Cloud-Clone Corp. | PAA068Ra02 |
| Anti-alpha-synuclein pS129, EP1536Y (rabbit) | 1/1000 | Abcam | ab51253 |
| Antibody anti-mouse – Alexa488 (goat) | 1/500 | Jackson ImmunoResearch | 115-545-003 |
| Antibody anti-rabbit –Alexa488 (donkey) | 1/500 | Jackson ImmunoResearch | 711-545-152 |
| Antibody anti-mouse –Cy3 (donkey) | 1/500 | Jackson ImmunoResearch | 715-165-150 |
| Antibody anti-rat – Cy5 (donkey) | 1/500 | Jackson ImmunoResearch | 712-605-153 |
| Antibody anti-rabbit biotinylated | 1/500 | Jackson ImmunoResearch | 711-064-152 |
| ABC-HRP complex |  | Vector Laboratories | PK-6100 |
| DAPI |  | Invitrogen | D1306 |
| ProLong Diamond Antifade Mountant |  | Invitrogen | P36970 |
| Natural PVA (print core: BB 0.4; layer height: 0.15 mm; print temp: 220_C, bed: 60_C; infill: 20%; build plate adhesion: 3 mm brim) |  | Ultimaker | 1528692-5K |

OCT, optimal cutting temperature; NDS, normal donkey serum; NGS, normal goat serum; H<sub>2</sub>O<sub>2</sub>, hydrogen peroxide; ABC-HRP, avidin-biotin - horseradish peroxidase; DAPI, 4',6-diamidino-2-phenylindole (DAPI)

## List of abbreviations

CD31: Cluster of differentiation 31
CSA: Cross-sectional areas
DAPI: 4’,6-diamidino-2-phenylindole
GLUT1: Glucose transporter 1
GM: Grey matter
HbO: Oxygenated hemoglobin
HbR: deoxygenated hemoglobin
MRI: Magnetic resonance imaging
NTL: Non-transgenic littermate
PD: Parkinson’s disease
pS129-α-syn: Phospho-α-syn (p-α-syn) at Ser129 site
sO_2_: Oxygen saturation
SVOT: Spiral volumetric optoacoustic tomography
T1w: T1-weighted
WM: White matter
α-syn: Alpha-synuclein

## References

2018. Global, regional, and national burden of Parkinson’s disease, 1990-2016: a systematic analysis for the Global Burden of Disease Study 2016. Lancet Neurol 17(11), 939-953.

American National Standards, I., Laser Institute of, A., 2022. ANSI Z136.1 Safe Use of Lasers - 2022. Laser Institute of America.

Anderson, J.P., Walker, D.E., Goldstein, J.M., de Laat, R., Banducci, K., Caccavello, R.J., Barbour, R., Huang, J., Kling, K., Lee, M., Diep, L., Keim, P.S., Shen, X., Chataway, T., Schlossmacher, M.G., Seubert, P., Schenk, D., Sinha, S., Gai, W.P., Chilcote, T.J., 2006. Phosphorylation of Ser-129 is the dominant pathological modification of alpha-synuclein in familial and sporadic Lewy body disease. J Biol Chem 281(40), 29739–29752.

Benskey, M.J., Perez, R.G., Manfredsson, F.P., 2016. The contribution of alpha synuclein to neuronal survival and function - Implications for Parkinson’s disease. J Neurochem 137(3), 331–359.

Bilgen, M., Al-Hafez, B., Berman, N.E., Festoff, B.W., 2005. Magnetic resonance imaging of mouse spinal cord. Magn Reson Med 54(5), 1226–1231.

Blauwendraat, C., Nalls, M.A., Singleton, A.B., 2020. The genetic architecture of Parkinson’s disease. Lancet Neurol 19(2), 170–178.

Braak, H., Del Tredici, K., 2017. Neuropathological Staging of Brain Pathology in Sporadic Parkinson’s disease: Separating the Wheat from the Chaff. J Parkinsons Dis 7(s1), S71–s85.

Braak, H., Del Tredici, K., Rüb, U., de Vos, R.A., Jansen Steur, E.N., Braak, E., 2003. Staging of brain pathology related to sporadic Parkinson’s disease. Neurobiol Aging 24(2), 197–211.

Bukkuri, A., Gatenby, R.A., Brown, J.S., 2022. GLUT1 production in cancer cells: a tragedy of the commons. NPJ Syst Biol Appl 8(1), 22.

Burtscher, J., Duderstadt, Y., Gatterer, H., Burtscher, M., Vozdek, R., Millet, G.P., Hicks, A.A., Ehrenreich, H., Kopp, M., 2024. Hypoxia Sensing and Responses in Parkinson’s Disease. Int J Mol Sci 25(3).

Butovsky, O., Weiner, H.L., 2018. Microglial signatures and their role in health and disease. Nat Rev Neurosci 19(10), 622–635.

Chen, T., Li, J., Chao, D., Sandhu, H.K., Liao, X., Zhao, J., Wen, G., Xia, Y., 2014. δ-Opioid receptor activation reduces α-synuclein overexpression and oligomer formation induced by MPP(+) and/or hypoxia. Exp Neurol 255, 127–136.

Chen, Z., Zhou, Q., Deán-Ben, X.L., Gezginer, I., Ni, R., Reiss, M., Shoham, S., Razansky, D., 2022. Multimodal Noninvasive Functional Neurophotonic Imaging of Murine Brain-Wide Sensory Responses. Adv Sci (Weinh) 9(24), e2105588.

Chera, B., Schaecher, K.E., Rocchini, A., Imam, S.Z., Ray, S.K., Ali, S.F., Banik, N.L., 2002. Calpain upregulation and neuron death in spinal cord of MPTP-induced parkinsonism in mice. Ann N Y Acad Sci 965, 274–280.

Chera, B., Schaecher, K.E., Rocchini, A., Imam, S.Z., Sribnick, E.A., Ray, S.K., Ali, S.F., Banik, N.L., 2004. Immunofluorescent labeling of increased calpain expression and neuronal death in the spinal cord of 1-methyl-4-phenyl-1,2,3,6-tetrahydropyridine-treated mice. Brain Res 1006(2), 150-156.

Chu, W.T., DeSimone, J.C., Riffe, C.J., Liu, H., Chakrabarty, P., Giasson, B.I., Vedam-Mai, V., Vaillancourt, D.E., 2020. α-Synuclein Induces Progressive Changes in Brain Microstructure and Sensory-Evoked Brain Function That Precedes Locomotor Decline. J Neurosci 40(34), 6649–6659.

Claron, J., Hingot, V., Rivals, I., Rahal, L., Couture, O., Deffieux, T., Tanter, M., Pezet, S., 2021. Large-scale functional ultrasound imaging of the spinal cord reveals in-depth spatiotemporal responses of spinal nociceptive circuits in both normal and inflammatory states. Pain 162(4), 1047-1059.

Cox, B., Laufer, J.G., Arridge, S.R., Beard, P.C., 2012. Quantitative spectroscopic photoacoustic imaging: a review. J Biomed Opt 17(6), 061202.

Davies, A.L., Desai, R.A., Bloomfield, P.S., McIntosh, P.R., Chapple, K.J., Linington, C., Fairless, R., Diem, R., Kasti, M., Murphy, M.P., Smith, K.J., 2013. Neurological deficits caused by tissue hypoxia in neuroinflammatory disease. Ann Neurol 74(6), 815–825.

De Leener, B., Lévy, S., Dupont, S.M., Fonov, V.S., Stikov, N., Louis Collins, D., Callot, V., Cohen-Adad, J., 2017. SCT: Spinal Cord Toolbox, an open-source software for processing spinal cord MRI data. Neuroimage 145(Pt A), 24–43.

Del Tredici, K., Braak, H., 2012. Spinal cord lesions in sporadic Parkinson’s disease. Acta Neuropathol 124(5), 643–664.

Delic, V., Chandra, S., Abdelmotilib, H., Maltbie, T., Wang, S., Kem, D., Scott, H.J., Underwood, R.N., Liu, Z., Volpicelli-Daley, L.A., West, A.B., 2018. Sensitivity and specificity of phospho-Ser129 α-synuclein monoclonal antibodies. J Comp Neurol 526(12), 1978–1990.

Deán-Ben, X.L., Robin, J., Nozdriukhin, D., Ni, R., Zhao, J., Glück, C., Droux, J., Sendón-Lago, J., Chen, Z., Zhou, Q., Weber, B., Wegener, S., Vidal, A., Arand, M., El Amki, M., Razansky, D., 2023. Deep optoacoustic localization microangiography of ischemic stroke in mice. Nat Commun 14(1), 3584.

Elfarrash, S., Jensen, N.M., Ferreira, N., Schmidt, S.I., Gregersen, E., Vestergaard, M.V., Nabavi, S., Meyer, M., Jensen, P.H., 2021. Polo-like kinase 2 inhibition reduces serine-129 phosphorylation of physiological nuclear alpha-synuclein but not of the aggregated alpha-synuclein. PLoS One 16(10), e0252635.

Esipova, T.V., Barrett, M.J.P., Erlebach, E., Masunov, A.E., Weber, B., Vinogradov, S.A., 2019. Oxyphor 2P: A High-Performance Probe for Deep-Tissue Longitudinal Oxygen Imaging. Cell Metab 29(3), 736–744.e737.

Fiederling, F., Hammond, L.A., Ng, D., Mason, C., Dodd, J., 2021. SpineRacks and SpinalJ for efficient analysis of neurons in a 3D reference atlas of the mouse spinal cord. STAR Protoc 2(4), 100897.

Finkelstein, D.I., Hare, D.J., Billings, J.L., Sedjahtera, A., Nurjono, M., Arthofer, E., George, S., Culvenor, J.G., Bush, A.I., Adlard, P.A., 2016. Clioquinol Improves Cognitive, Motor Function, and Microanatomy of the Alpha-Synuclein hA53T Transgenic Mice. ACS Chem Neurosci 7(1), 119-129.

Fuentes, R., Petersson, P., Siesser, W.B., Caron, M.G., Nicolelis, M.A., 2009. Spinal cord stimulation restores locomotion in animal models of Parkinson’s disease. Science 323(5921), 1578–1582.

Fujiwara, H., Hasegawa, M., Dohmae, N., Kawashima, A., Masliah, E., Goldberg, M.S., Shen, J., Takio, K., Iwatsubo, T., 2002. alpha-Synuclein is phosphorylated in synucleinopathy lesions. Nat Cell Biol 4(2), 160-164.

Gao, J., Jiang, M., Magin, R.L., Gatto, R.G., Morfini, G., Larson, A.C., Li, W., 2020. Multicomponent diffusion analysis reveals microstructural alterations in spinal cord of a mouse model of amyotrophic lateral sclerosis ex vivo. PLoS One 15(4), e0231598.

Garcia-Esparcia, P., Hernández-Ortega, K., Koneti, A., Gil, L., Delgado-Morales, R., Castaño, E., Carmona, M., Ferrer, I., 2015. Altered machinery of protein synthesis is region-and stage-dependent and is associated with α-synuclein oligomers in Parkinson’s disease. Acta Neuropathol Commun 3, 76.

Gatto, R.G., Li, W., Gao, J., Magin, R.L., 2018. In vivo diffusion MRI detects early spinal cord axonal pathology in a mouse model of amyotrophic lateral sclerosis. NMR Biomed 31(8), e3954.

Giasson, B.I., Duda, J.E., Quinn, S.M., Zhang, B., Trojanowski, J.Q., Lee, V.M., 2002. Neuronal alpha-synucleinopathy with severe movement disorder in mice expressing A53T human alpha-synuclein. Neuron 34(4), 521–533.

Goedert, M., Clavaguera, F., Tolnay, M., 2010. The propagation of prion-like protein inclusions in neurodegenerative diseases. Trends Neurosci 33(7), 317–325.

Gottschalk, S., Degtyaruk, O., Mc Larney, B., Rebling, J., Hutter, M.A., Deán-Ben, X.L., Shoham, S., Razansky, D., 2019. Rapid volumetric optoacoustic imaging of neural dynamics across the mouse brain. Nature Biomedical Engineering 3(5), 392–401.

Guo, M., Ji, X., Liu, J., 2022. Hypoxia and Alpha-Synuclein: Inextricable Link Underlying the Pathologic Progression of Parkinson’s Disease. Front Aging Neurosci 14, 919343.

Guo, M., Liu, W., Luo, H., Shao, Q., Li, Y., Gu, Y., Guan, Y., Ma, W., Chen, M., Yang, H., Ji, X., Liu, J., 2023. Hypoxic stress accelerates the propagation of pathological alpha-synuclein and degeneration of dopaminergic neurons. CNS Neurosci Ther 29(2), 544–558.

Harmon, J.N., Chandran, P., Chandrasekaran, A., Hyde, J.E., Hernandez, G.J., Reed, M.J., Bruce, M.F., Khaing, Z.Z., 2024. Contrast-enhanced ultrasound imaging detects anatomical and functional changes in rat cervical spine microvasculature with normal aging. bioRxiv.

Hatcher, J.M., Pennell, K.D., Miller, G.W., 2008. Parkinson’s disease and pesticides: a toxicological perspective. Trends Pharmacol Sci 29(6), 322–329.

Hernandez-Gerez, E., Fleming, I.N., Parson, S.H., 2019. A role for spinal cord hypoxia in neurodegeneration. Cell Death Dis 10(11), 861.

Hijaz, B.A., Volpicelli-Daley, L.A., 2020. Initiation and propagation of α-synuclein aggregation in the nervous system. Molecular Neurodegeneration 15(1), 19.

Hochuli, R., An, L., Beard, P.C., Cox, B.T., 2019. Estimating blood oxygenation from photoacoustic images: can a simple linear spectroscopic inversion ever work? J Biomed Opt 24(12), 1–13.

Huang, Z., Xu, Z., Wu, Y., Zhou, Y., 2011. Determining nuclear localization of alpha-synuclein in mouse brains. Neuroscience 199, 318–332.

Höglinger, G.U., Adler, C.H., Berg, D., Klein, C., Outeiro, T.F., Poewe, W., Postuma, R., Stoessl, A.J., Lang, A.E., 2024. A biological classification of Parkinson’s disease: the SynNeurGe research diagnostic criteria. Lancet Neurol 23(2), 191–204.

Isensee, F., Jaeger, P.F., Kohl, S.A.A., Petersen, J., Maier-Hein, K.H., 2021. nnU-Net: a self-configuring method for deep learning-based biomedical image segmentation. Nat Methods 18(2), 203–211.

Jacques, S.L., 2013. Optical properties of biological tissues: a review. Phys Med Biol 58(11), R37-61.

Kalva, S.K., Dean-Ben, X.L., Razansky, D., 2021. Single-sweep volumetric optoacoustic tomography of whole mice. Photonics Research 9(6), 899-908.

Kalva, S.K., Deán-Ben, X.L., Reiss, M., Razansky, D., 2023a. Head-to-tail imaging of mice with spiral volumetric optoacoustic tomography. Photoacoustics 30, 100480.

Kalva, S.K., Deán-Ben, X.L., Reiss, M., Razansky, D., 2023b. Spiral volumetric optoacoustic tomography for imaging whole-body biodynamics in small animals. Nat Protoc 18(7), 2124–2142.

Kalva, S.K., Sánchez-Iglesias, A., Deán-Ben, X.L., Liz-Marzán, L.M., Razansky, D., 2022. Rapid Volumetric Optoacoustic Tracking of Nanoparticle Kinetics across Murine Organs. ACS Appl Mater Interfaces 14(1), 172–178.

Kecheliev, V., Boss, L., Maheshwari, U., Konietzko, U., Keller, A., Razansky, D., Nitsch, R.M., Klohs, J., Ni, R., 2023. Aquaporin 4 is differentially increased and dislocated in association with tau and amyloid-beta. Life Sciences, 121593.

Kim, T., Mehta, S.L., Kaimal, B., Lyons, K., Dempsey, R.J., Vemuganti, R., 2016. Poststroke Induction of α-Synuclein Mediates Ischemic Brain Damage. J Neurosci 36(26), 7055–7065.

Kong, Y., Maschio, C.A., Shi, X., Xie, F., Zuo, C., Konietzko, U., Shi, K., Rominger, A., Xiao, J., Huang, Q., Nitsch, R.M., Guan, Y., Ni, R., 2024. Relationship Between Reactive Astrocytes, by [(18)F]SMBT-1 Imaging, with Amyloid-Beta, Tau, Glucose Metabolism, and TSPO in Mouse Models of Alzheimer’s Disease. Mol Neurobiol.

Kontopoulos, E., Parvin, J.D., Feany, M.B., 2006. Alpha-synuclein acts in the nucleus to inhibit histone acetylation and promote neurotoxicity. Hum Mol Genet 15(20), 3012–3023.

Kumar, S.T., Jagannath, S., Francois, C., Vanderstichele, H., Stoops, E., Lashuel, H.A., 2020. How specific are the conformation-specific α-synuclein antibodies? Characterization and validation of 16 α-synuclein conformation-specific antibodies using well-characterized preparations of α-synuclein monomers, fibrils and oligomers with distinct structures and morphology. Neurobiol Dis 146, 105086.

Laakso, H., Lehto, L.J., Paasonen, J., Salo, R., Canna, A., Lavrov, I., Michaeli, S., Gröhn, O., Mangia, S., 2021. Spinal cord fMRI with MB-SWIFT for assessing epidural spinal cord stimulation in rats. Magn Reson Med 86(4), 2137-2145.

Landelle, C., Dahlberg, L.S., Lungu, O., Misic, B., De Leener, B., Doyon, J., 2023. Altered Spinal Cord Functional Connectivity Associated with Parkinson’s Disease Progression. Mov Disord 38(4), 636–645.

Lashuel, H.A., Mahul-Mellier, A.L., Novello, S., Hegde, R.N., Jasiqi, Y., Altay, M.F., Donzelli, S., DeGuire, S.M., Burai, R., Magalhães, P., Chiki, A., Ricci, J., Boussouf, M., Sadek, A., Stoops, E., Iseli, C., Guex, N., 2022. Revisiting the specificity and ability of phospho-S129 antibodies to capture alpha-synuclein biochemical and pathological diversity. NPJ Parkinsons Dis 8(1), 136.

Lestón Pinilla, L., Ugun-Klusek, A., Rutella, S., De Girolamo, L.A., 2021. Hypoxia Signaling in Parkinson’s Disease: There Is Use in Asking “What HIF?”. Biology (Basel) 10(8).

Li, G., Liu, J., Guo, M., Gu, Y., Guan, Y., Shao, Q., Ma, W., Ji, X., 2022. Chronic hypoxia leads to cognitive impairment by promoting HIF-2α-mediated ceramide catabolism and alpha-synuclein hyperphosphorylation. Cell Death Discovery 8(1), 473.

Liu, C., Cárdenas-Rivera, A., Teitelbaum, S., Birmingham, A., Alfadhel, M., Yaseen, M.A., 2024. Neuroinflammation increases oxygen extraction in a mouse model of Alzheimer’s disease. Alzheimers Res Ther 16(1), 78.

Massalimova, A., Ni, R., Nitsch, R.M., Reisert, M., von Elverfeldt, D., Klohs, J., 2021. Diffusion Tensor Imaging Reveals Whole-Brain Microstructural Changes in the P301L Mouse Model of Tauopathy. Neurodegener Dis, 1-12.

Matsubayashi, K., Nagoshi, N., Komaki, Y., Kojima, K., Shinozaki, M., Tsuji, O., Iwanami, A., Ishihara, R., Takata, N., Matsumoto, M., Mimura, M., Okano, H., Nakamura, M., 2018. Assessing cortical plasticity after spinal cord injury by using resting-state functional magnetic resonance imaging in awake adult mice. Sci Rep 8(1), 14406.

Melzer, T.R., Watts, R., MacAskill, M.R., Pearson, J.F., Rüeger, S., Pitcher, T.L., Livingston, L., Graham, C., Keenan, R., Shankaranarayanan, A., Alsop, D.C., Dalrymple-Alford, J.C., Anderson, T.J., 2011. Arterial spin labelling reveals an abnormal cerebral perfusion pattern in Parkinson’s disease. Brain 134(Pt 3), 845–855.

Meyer, B.P., Hirschler, L., Lee, S., Kurpad, S.N., Warnking, J.M., Barbier, E.L., Budde, M.D., 2021. Optimized cervical spinal cord perfusion MRI after traumatic injury in the rat. J Cereb Blood Flow Metab 41(8), 2010–2025.

Miller, R.M., Kiser, G.L., Kaysser-Kranich, T., Casaceli, C., Colla, E., Lee, M.K., Palaniappan, C., Federoff, H.J., 2007. Wild-type and mutant alpha-synuclein induce a multi-component gene expression profile consistent with shared pathophysiology in different transgenic mouse models of PD. Exp Neurol 204(1), 421–432.

Mondal, R., Campoy, A.-D.T., Liang, C., Mukherjee, J., 2021. [18F]FDG PET/CT Studies in Transgenic Hualpha-Syn (A53T) Parkinson’s Disease Mouse Model of α-Synucleinopathy. Frontiers in Neuroscience 15, 718.

Murata, H., Barnhill, L.M., Bronstein, J.M., 2022. Air Pollution and the Risk of Parkinson’s Disease: A Review. Mov Disord 37(5), 894–904.

Ni, R., 2022. PET imaging in animal models of Parkinson’s disease. Behavioural Brain Research, 114174.

Ni, R., Chen, Z., Deán-Ben, X.L., Voigt, F.F., Kirschenbaum, D., Shi, G., Villois, A., Zhou, Q., Crimi, A., Arosio, P., Nitsch, R.M., Nilsson, K.P.R., Aguzzi, A., Helmchen, F., Klohs, J., Razansky, D., 2022. Multiscale optical and optoacoustic imaging of amyloid-β deposits in mice. Nat Biomed Eng 6(9), 1031–1044.

Ni, R., Rudin, M., Klohs, J., 2018. Cortical hypoperfusion and reduced cerebral metabolic rate of oxygen in the arcAβ mouse model of Alzheimer’s disease. Photoacoustics 10, 38–47.

Ni, R., Straumann, N., Fazio, S., Dean-Ben, X.L., Louloudis, G., Keller, C., Razansky, D., Ametamey, S., Mu, L., Nombela-Arrieta, C., Klohs, J., 2023. Imaging increased metabolism in the spinal cord in mice after middle cerebral artery occlusion. Photoacoustics 32, 100532.

Ni, R., Zarb, Y., Kuhn, G.A., Müller, R., Yundung, Y., Nitsch, R.M., Kulic, L., Keller, A., Klohs, J., 2020. SWI and phase imaging reveal intracranial calcifications in the P301L mouse model of human tauopathy. Magma 33(6), 769–781.

Norris, E.H., Uryu, K., Leight, S., Giasson, B.I., Trojanowski, J.Q., Lee, V.M., 2007. Pesticide exposure exacerbates alpha-synucleinopathy in an A53T transgenic mouse model. Am J Pathol 170(2), 658–666.

Outeiro, T.F., Putcha, P., Tetzlaff, J.E., Spoelgen, R., Koker, M., Carvalho, F., Hyman, B.T., McLean, P.J., 2008. Formation of toxic oligomeric alpha-synuclein species in living cells. PLoS One 3(4), e1867.

Pang, S.Y.-Y., Ho, P.W.-L., Liu, H.-F., Leung, C.-T., Li, L., Chang, E.E.S., Ramsden, D.B., Ho, S.-L., 2019. The interplay of aging, genetics and environmental factors in the pathogenesis of Parkinson’s disease. Translational Neurodegeneration 8(1), 23.

Parra-Rivas, L.A., Madhivanan, K., Aulston, B.D., Wang, L., Prakashchand, D.D., Boyer, N.P., Saia-Cereda, V.M., Branes-Guerrero, K., Pizzo, D.P., Bagchi, P., Sundar, V.S., Tang, Y., Das, U., Scott, D.A., Rangamani, P., Ogawa, Y., Subhojit, R., 2023. Serine-129 phosphorylation of α-synuclein is an activity-dependent trigger for physiologic protein-protein interactions and synaptic function. Neuron 111(24), 4006–4023.e4010.

Pinho, R., Paiva, I., Jercic, K.G., Fonseca-Ornelas, L., Gerhardt, E., Fahlbusch, C., Garcia-Esparcia, P., Kerimoglu, C., Pavlou, M.A.S., Villar-Piqué, A., Szego, É., Lopes da Fonseca, T., Odoardi, F., Soeroes, S., Rego, A.C., Fischle, W., Schwamborn, J.C., Meyer, T., Kügler, S., Ferrer, I., Attems, J., Fischer, A., Becker, S., Zweckstetter, M., Borovecki, F., Outeiro, T.F., 2019. Nuclear localization and phosphorylation modulate pathological effects of alpha-synuclein. Hum Mol Genet 28(1), 31–50.

Poewe, W., Seppi, K., Tanner, C.M., Halliday, G.M., Brundin, P., Volkmann, J., Schrag, A.-E., Lang, A.E., 2017. Parkinson disease. Nature Reviews Disease Primers 3(1), 17013.

Ramalingam, N., Jin, S.X., Moors, T.E., Fonseca-Ornelas, L., Shimanaka, K., Lei, S., Cam, H.P., Watson, A.H., Brontesi, L., Ding, L., Hacibaloglu, D.Y., Jiang, H., Choi, S.J., Kanter, E., Liu, L., Bartels, T., Nuber, S., Sulzer, D., Mosharov, E.V., Chen, W.V., Li, S., Selkoe, D.J., Dettmer, U., 2023. Dynamic physiological α-synuclein S129 phosphorylation is driven by neuronal activity. NPJ Parkinsons Dis 9(1), 4.

Ramos-Vega, M., Kjellman, P., Todorov, M.I., Kylkilahti, T.M., Bäckström, B.T., Ertürk, A., Madsen, C.D., Lundgaard, I., 2022. Mapping of neuroinflammation-induced hypoxia in the spinal cord using optoacoustic imaging. Acta Neuropathol Commun 10(1), 51.

Ron, A., Deán-Ben, X.L., Gottschalk, S., Razansky, D., 2019a. Volumetric Optoacoustic Imaging Unveils High-Resolution Patterns of Acute and Cyclic Hypoxia in a Murine Model of Breast Cancer. Cancer Res 79(18), 4767–4775.

Ron, A., Deán-Ben, X.L., Reber, J., Ntziachristos, V., Razansky, D., 2019b. Characterization of Brown Adipose Tissue in a Diabetic Mouse Model with Spiral Volumetric Optoacoustic Tomography. Mol Imaging Biol 21(4), 620–625.

Rust, R., Grönnert, L., Gantner, C., Enzler, A., Mulders, G., Weber, R.Z., Siewert, A., Limasale, Y.D.P., Meinhardt, A., Maurer, M.A., Sartori, A.M., Hofer, A.S., Werner, C., Schwab, M.E., 2019. Nogo-A targeted therapy promotes vascular repair and functional recovery following stroke. Proc Natl Acad Sci U S A 116(28), 14270–14279.

Saito, S., Mori, Y., Yoshioka, Y., Murase, K., 2012. High-resolution ex vivo imaging in mouse spinal cord using micro-CT with 11.7T-MRI and myelin staining validation. Neuroscience Research 73(4), 337–340.

Samantaray, S., Knaryan, V.H., Guyton, M.K., Matzelle, D.D., Ray, S.K., Banik, N.L., 2007. The parkinsonian neurotoxin rotenone activates calpain and caspase-3 leading to motoneuron degeneration in spinal cord of Lewis rats. Neuroscience 146(2), 741–755.

Sartoretti, T., Ganley, R.P., Ni, R., Freund, P., Zeilhofer, H.U., Klohs, J., 2022. Structural MRI Reveals Cervical Spinal Cord Atrophy in the P301L Mouse Model of Tauopathy: Gender and Transgene-Dosing Effects. Frontiers in Aging Neuroscience 14.

Schell, H., Hasegawa, T., Neumann, M., Kahle, P.J., 2009. Nuclear and neuritic distribution of serine-129 phosphorylated alpha-synuclein in transgenic mice. Neuroscience 160(4), 796–804.

Shibata, M., Ohtani, R., Ihara, M., Tomimoto, H., 2004. White matter lesions and glial activation in a novel mouse model of chronic cerebral hypoperfusion. Stroke 35(11), 2598–2603.

Siddiqui, A., Chinta, S.J., Mallajosyula, J.K., Rajagopolan, S., Hanson, I., Rane, A., Melov, S., Andersen, J.K., 2012. Selective binding of nuclear alpha-synuclein to the PGC1alpha promoter under conditions of oxidative stress may contribute to losses in mitochondrial function: implications for Parkinson’s disease. Free Radic Biol Med 53(4), 993–1003.

Simuni, T., Chahine, L.M., Poston, K., Brumm, M., Buracchio, T., Campbell, M., Chowdhury, S., Coffey, C., Concha-Marambio, L., Dam, T., DiBiaso, P., Foroud, T., Frasier, M., Gochanour, C., Jennings, D., Kieburtz, K., Kopil, C.M., Merchant, K., Mollenhauer, B., Montine, T., Nudelman, K., Pagano, G., Seibyl, J., Sherer, T., Singleton, A., Stephenson, D., Stern, M., Soto, C., Tanner, C.M., Tolosa, E., Weintraub, D., Xiao, Y., Siderowf, A., Dunn, B., Marek, K., 2024. A biological definition of neuronal α-synuclein disease: towards an integrated staging system for research. Lancet Neurol 23(2), 178–190.

Sobek, J., Li, J., Combes, B.F., Gerez, J.A., Nilsson, P.K., Henrich, M.T., Geibl, F.F., Shi, K., Rominger, A., Oertel, W.H., 2023. Efficient characterization of multiple binding sites of small molecule imaging ligands on amyloid-beta, 4-repeat/full-length tau and alpha-synuclein. bioRxiv, 2023-2003.

Sorrentino, Z.A., Xia, Y., Funk, C., Riffe, C.J., Rutherford, N.J., Ceballos Diaz, C., Sacino, A.N., Price, N.D., Golde, T.E., Giasson, B.I., Chakrabarty, P., 2018. Motor neuron loss and neuroinflammation in a model of α-synuclein-induced neurodegeneration. Neurobiol Dis 120, 98-106.

Straumann, N., Combes, B.F., Dean-Ben, X.L., Steernke, R., Gerez, J., Dias, I., Chen, Z., Watts, B., Rostami, I., Shi, K., 2023. Visualizing alpha-synuclein and iron deposition in M83 mouse model of Parkinson’s disease in vivo. bioRxiv, 2023-2006.

Sun, H.L., Sun, B.L., Chen, D.W., Chen, Y., Li, W.W., Xu, M.Y., Shen, Y.Y., Xu, Z.Q., Wang, Y.J., Bu, X.L., 2019. Plasma α-synuclein levels are increased in patients with obstructive sleep apnea syndrome. Ann Clin Transl Neurol 6(4), 788–794.

Unal-Cevik, I., Gursoy-Ozdemir, Y., Yemisci, M., Lule, S., Gurer, G., Can, A., Müller, V., Kahle, P.J., Dalkara, T., 2011. Alpha-synuclein aggregation induced by brief ischemia negatively impacts neuronal survival in vivo: a study in [A30P]alpha-synuclein transgenic mouse. J Cereb Blood Flow Metab 31(3), 913–923.

Vagenknecht, P., Luzgin, A., Ono, M., Ji, B., Higuchi, M., Noain, D., Maschio, C.A., Sobek, J., Chen, Z., Konietzko, U., Gerez, J.A., Riek, R., Razansky, D., Klohs, J., Nitsch, R.M., Dean-Ben, X.L., Ni, R., 2022. Non-invasive imaging of tau-targeted probe uptake by whole brain multi-spectral optoacoustic tomography. Eur J Nucl Med Mol Imaging 49(7), 2137–2152.

van Horssen, J., van Schaik, P., Witte, M., 2019. Inflammation and mitochondrial dysfunction: A vicious circle in neurodegenerative disorders? Neurosci Lett 710, 132931.

Watson, C., Paxinos, G., Kayalioglu, G., Heise, C., 2009. Chapter 16 - Atlas of the Mouse Spinal Cord. In: Watson, C., Paxinos, G., Kayalioglu, G. (Eds.), The Spinal Cord, pp. 308–379. Academic Press, San Diego.

Wolters, F.J., Zonneveld, H.I., Hofman, A., van der Lugt, A., Koudstaal, P.J., Vernooij, M.W., Ikram, M.A., 2017. Cerebral Perfusion and the Risk of Dementia: A Population-Based Study. Circulation 136(8), 719–728.

Wu, W., He, S., Wu, J., Chen, C., Li, X., Liu, K., Qu, J.Y., 2022. Long-term in vivo imaging of mouse spinal cord through an optically cleared intervertebral window. Nature Communications 13(1), 1959.

Yang, R., Dunn, J.F., 2019. Multiple sclerosis disease progression: Contributions from a hypoxia-inflammation cycle. Mult Scler 25(13), 1715-1718.

